# Gastrula-premarked posterior enhancer primes posterior tissue development through cross-talk with TGF-β signaling pathway

**DOI:** 10.1101/2024.04.14.589453

**Authors:** Yingying Chen, Fengxiang Tan, Qing Fang, Lin Zhang, Jiaoyang Liao, Penglei Shen, Yun Qian, Mingzhu Wen, Rui Song, Yonggao Fu, He Jax Xu, Ran Wang, Cheng Li, Zhen Shao, Jinsong Li, Naihe Jing, Xianfa Yang

## Abstract

The regulatory mechanisms governing cell fate determination, particularly lineage diversification during mammalian embryonic development, remain poorly understood with in-depth regulatory paradigms yet to be fully elucidated. Here, leveraging the epigenetic landscape of mouse gastrula, we identified p-Enh, a pre-marked enhancer in primitive streak region, as pivotal regulator for posterior tissue development in mouse embryos. Morphological and single-cell transcriptomic analyses confirmed embryonic lethality phenotype with disrupted posterior tissue development trajectories in p-Enh-KO embryos. Molecularly, apart from regulating the neighboring coding-gene *Cdx2 in cis*, our findings suggest that p-Enh also modulate the global transcriptome and epigenomic landscape, which might through the transient production of eRNA *in trans*. Further investigation revealed p-Enh participate in the regulatory cascades of TGF-β signaling. Chemical modulation of TGF-β signaling can largely rescue the posterior development deficiency in *in vitro* gastruloids through a *Cdx2*-independent mechanism. Thus, we propose a potential model in which the broadly distributed p-Enh transcripts within the nucleus could serve as essential cross-modular coordinators, priming the posterior development of mouse embryo.

## 1. Introduction

Gastrulation stands as a pivotal developmental process to set up the embryonic body plan, characterized by the differentiation of the epiblast into three germ layers ^[1,2]^. Studies reported that the posteriorly positioned primitive streak (PS) and adjacent posterior epiblast region can integrate the intricate developmental signals, and then serve as a progenitor reservoir with the competence to give rise to a diverse range of tissues, spanning from mesodermal tissues such as the anlage of somitic, limb and cardiac mesoderm to the posterior neural lineage of the spinal cord ^[3–5]^. Nonetheless, how cells reside in the PS region coordinate the extrinsic signals and intrinsic molecular features and sustain the competence for prospective cell fate differentiation remains largely elusive.

Vital developmental morphogens lay the foundation for tissue patterning and homeostasis across various organs ^[6]^. The cross-talk among different pathways forms a “signal cocktail” with an optimal concentration niche, which facilitates diverse cell differentiation programs. For instance, during gastrulation, Wnt and TGF-β signaling pathways are activated in the PS and mesodermal cells, then diffuse from the organized sources and form morphogen gradients, thus instructing the cell fate of surrounding tissues ^[7,8]^. However, β-catenin and SMAD4, the downstream effectors of Wnt and TGF-β signaling, are broadly expressed throughout the entire embryo during gastrulation ^[9]^. Distinct sets of downstream developmental genes are then activated to promote cell fate diversification in different regions of the embryo ^[1]^. The specific underlying mechanisms for how these signals anchor precisely to a specific set of developmental genes remain as mysteries.

The selective chromatin responsiveness, manifested as various chromatin accessibilities, chromatin modifications, or chromatin topologies at regulatory loci, to signal effectors and co-factors, have been reported as an important dimension in orchestrating the regulative forces of extrinsic signals ^[10]^. Notably, the regulatory elements are frequently docking platforms for transcription factors (TFs) and are usually annotated as enhancers or promoters ^[11]^. Dynamics of regulatory elements contribute to the regulation of lineage-specific transcriptional programs under distinct developmental conditions ^[12]^. Mechanistic studies indicate that regulatory elements can not only regulate the expression of neighboring genes but also hold the proper cell fate plasticity through direct enhancer-promoter chromatin interactions or self-transcribed enhancer RNAs (eRNAs) ^[13,14]^. In particular, the discovery of eRNAs has expanded the understanding of enhancer functions from modulating local gene targets *in cis* to even genome-wide chromatin remodeling and cross-modular interactions with the transcriptional machinery *in trans* ^[15–17]^. Recently, we and others have reported that enhancers crucial for prospective tissues development are frequently pre-marked by epigenetic modifications or TFs’ binding prior to the corresponding gene expression ^[13,18–22]^, a phenomena termed “chromatin priming”. However, limited knowledge has been acquired concerning the coordination between chromatin regulation and the extrinsic signals in the establishment of cellular developmental competence and the subsequent lineage diversification of the PS region.

In this study, through combined bioinformatic analysis and experimental screening of the pre-marked regulatory elements in the PS regions of the mouse gastrula, we identified a regulatory element, p-Enh, which is located in the first intron of *Cdx2.* Transgenic enhancer reporter assay revealed that p-Enh is pre-activated in the PS region at the late gastrulation stage (E7.5) and consistently activated in the tailbud region during early organogenesis. Phenotypic analyses of the p-Enh knockout embryo mutants revealed embryo lethality at around the E10.5 stage with severe disruptions of posterior tissue development. Single-cell RNA-seq confirmed that the cellular abundances of posterior tissues, such as neuromesodermal progenitors (NMP), limb mesenchyme, and somitic mesoderm, were heavily decreased. Mechanistically, besides regulating its neighboring coding-gene *Cdx2*, p-Enh could transiently express abundant eRNAs, in the nuclei of the PS cells at E7.5 but not in later-stage embryos. The genetic removal of p-Enh in *in vitro* differentiated NMP leads to dramatic remodeling of epigenomic landscape specifically at SMAD binding hotspots independent of CDX2 dependency. Furthermore, functional assays demonstrate that adjusting TGF-β signaling can rescue the p-Enh-KO defects. Thus, we propose that the transient expressed p-Enh-eRNAs might act as trans-modular coordinator, which bridges the specific interactions with morphogen effectors and specific genomic loci of master regulators, therefore contributing to the priming of posterior tissue development.

## 2. Results

### 2.1. Screening and identification of pre-marked distal regulatory elements within PS cells in the mouse gastrula

During mouse gastrulation, the PS cells firstly emerge at the E6.5 stage, then gradually form a morphologically distinct structure at the posterior embryonic region, and finally establish a diverse cell fate-specified progenitor pool at the late gastrulation stage (E7.0 to E7.5) ^[4,23]^. PS cells possess a remarkable capacity to give rise to a diverse range of tissues, encompassing mesodermal cells such as somites ^[24]^, limb mesenchyme ^[25]^, and cardiac mesoderm (Figure 1a) ^[26]^. Moreover, it has been recognized that the PS region of the mouse gastrula also harbors a newly identified bi-potent NMP, which can contribute to both the posterior spinal cord and mesoderm (Figure 1a) ^[27–29]^. Thus, the mechanisms governing PS cell development are fundamental for subsequent patterning of diverse tissues and organs.

**Figure 1.**
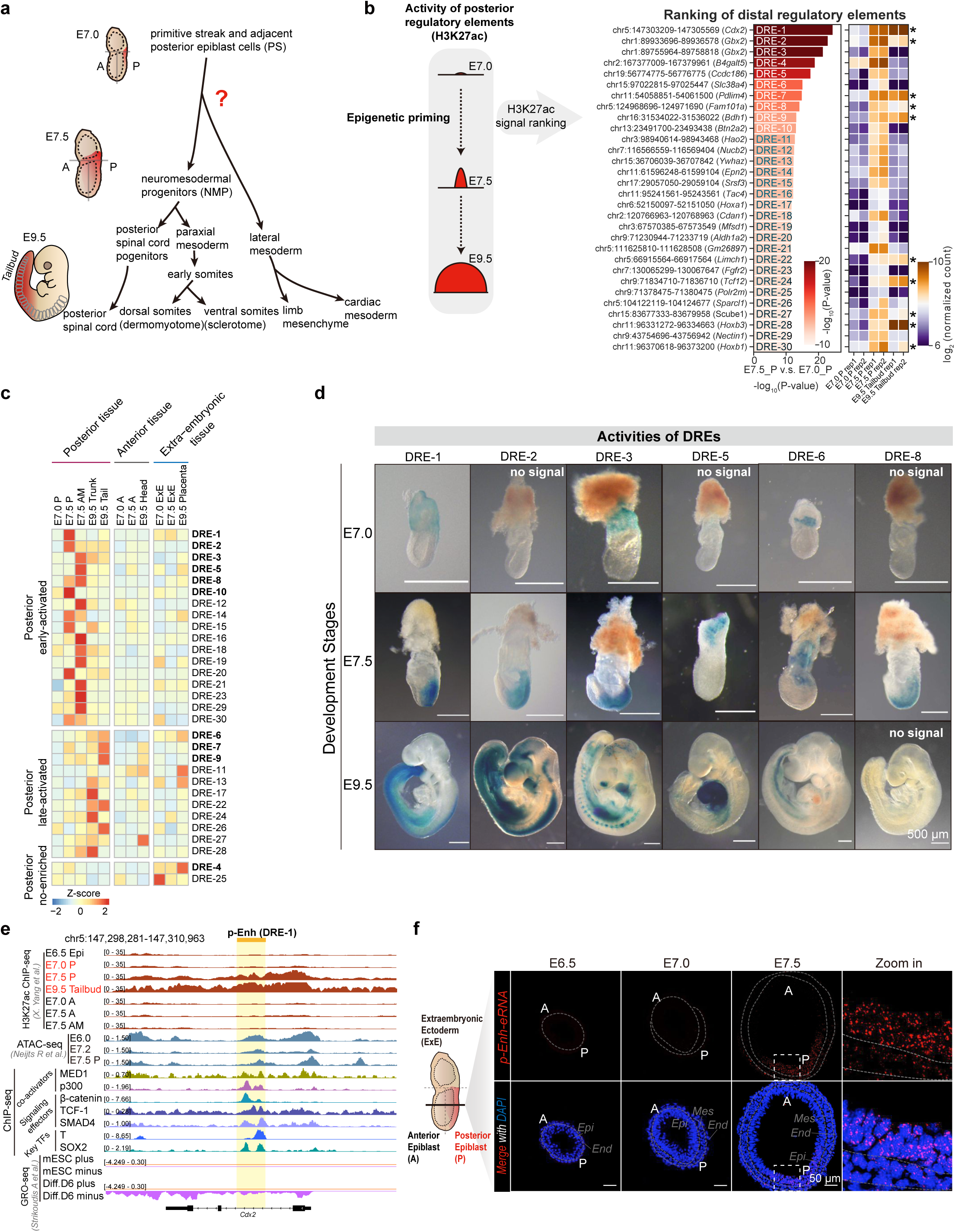
Screening and identification of pre-marked distal regulatory elements within PS cells in the mouse gastrula. **(a)** The diagram illustrating the lineages of posterior tissue development during mouse embryonic development. The precursor cells contributing to the posterior tissue are denoted in red shade. **(b)** The ranking of distal regulatory elements (DREs) and corresponding heatmap based on quantified H3K27ac ChIP-seq data ^[18,31]^. The top-30 significant DREs were shown based on calculated -log_10_(P-value). Heatmap was constructed based on log_2_(normalized count). DREs with retained abundant H3K37ac in E9.5 were marked with asterisk. **(c)** The heatmap visualizing the eRNA qPCR results correlated with Figure. 1b. Multiple tissue samples obtained from embryos at E7.0, E7.5, and E9.5. Tissue samples are stratified into three groups based on developmental trajectories. For each DRE locus, dual targeting segments are subjected for qPCR test, all tests were replicated in pairs to ensure reliability. The DREs highlighted in bold indicated DREs subjected to transgenic assay. **(d)** The LacZ staining revealing spatiotemporal activities of distinct DREs at key developmental stages (E7.0, E7.5, E9.5). (Some transgenic embryos with no detectable positive signals were labeled with ‘no signal’) Scale bar: 500 μm. **(e)** The genome browser snapshot showing chromatin status near p-Enh (DRE-1). The tracks including published H3K27ac ChIP-seq ^[18,31]^, ATAC-seq ^[111]^, ChIP-seq ^[38–44]^, and GRO-seq ^[45]^ datasets. **(f)** RNAscope targeting p-Enh-eRNA during gastrulation (E6.5, E7.0, E7.5). The schematic of the mouse gastrula is presented on the left side. Zoomed-in view showing the enrichment of p-Enh-eRNA at posterior epiblast. The labeling scheme employs “A” for anterior region, “P” for posterior region, “Epi” for epiblast, “Mes” for mesoderm, and “End” for endoderm. Scale bar: 50 μm.

Previously, we identified the epigenetic pre-mark at regulatory elements associated with developmental genes during gastrulation. This mechanism may serve as a widely available mechanism that primes prospective cell fate for following organogenesis ^[18]^. Here, to pinpoint the specific regulatory elements responsible for the developmental competence of the PS cells, we first employed MAnorm2 ^[30]^ to systematically analyze the dynamics of H3K27ac enrichment during the late gastrulation stage (from the E7.0 to the E7.5 stage) (Figure 1b). In total, 1,828 distal regulatory elements (DREs) were identified with significant upregulation (2-fold, p-value < 0.01) of H3K27ac enrichment within the posterior tissue (Figure S1a, Supporting Information). Genomic Regions Enrichment of Annotations Tool (GREAT) analysis of the top-100 DREs, ranked by p-value, revealed that these DREs are related to genes involved in anterior-posterior (A-P) patterning, animal organ morphogenesis, neural tube development, as well as mesenchymal cell differentiation, which further indicate that the pre-mark of H3K27ac at DREs are functionally related to the posterior tissue development (Figure S1b, Supporting Information; Table S1, Supporting Information). To determine the enrichment of H3K27ac during the following organogenesis, we checked the H3K27ac distribution at the DREs in E9.5 tailbud tissues by incorporating the published dataset ^[31]^. As shown in Figure 1b, among the top-30 ranked DREs, 10 DREs (marked with asterisk) maintain abundant H3K27ac enrichment in E9.5 tailbud.

Enhancer RNAs, which are transcribed from corresponding enhancers and often pre-loaded with RNA polymerase II and H3K27ac, could be used to represent the enhancer activity and also be involved in the regulation of gene expression and cell fate determination ^[32–36]^. To further refine the scope of potential functional DREs, we determined the abundance of eRNA transcripts for the top-30 DREs across various tissues of embryos at different stages (E7.0, E7.5, E9.5) using RT-qPCR (Figure 1c). Based on the expression pattern of these eRNAs, we clustered the eRNAs into three groups: posterior early-activated group with relatively high and specific eRNA expression in the PS cells of the E7.5 stage; posterior late-activated group with obvious eRNA expression in the E9.5 trunk and tailbud; and posterior no-enriched group (Figure 1c). Next, to determine the innate regulatory activity during mouse embryogenesis, we used *in vivo* transgenic enhancer reporter system to validate the DREs from the above three groups (Figure S1c, Supporting Information) ^[18,37]^. We performed transgenic reporter assays with the top-10 DREs, spanning the three eRNA groups (highlighted in bold in Figure 1c) and found that 6 out of 10 DREs could drive LacZ expression in the time window tested (E7.0 to E9.5). Notably, the majority (5 out of 6) of DREs with LacZ signals belonged to the early-activated group (DRE-1, 2, 3, 5, and 8). Only one DRE (DRE-6) from the late-activated group showed LacZ signal (1 out of 3), and no positive results were observed in the posterior no-enriched group (Figure 1d; Figure S1d, Supporting Information). Within these six DREs, DRE-1 exhibits the most pronounced caudal tissue specificity of LacZ signals. DRE-1 is *de novo* activated in the E7.5 PS cells with extraembryonic activity in E7.0 embryos, and its posterior region-specific enhancer activity is sustained in the tailbud region till later organogenetic stages (E8.5, E9.5 and E11.5) (Figure 1d; Figure S1e, Supporting Information). Further exploration of the genomic features of DRE-1 revealed that the evolutionarily conserved DRE-1 element is located in the first intron of the coding-gene, *Cdx2*, and harbors pervasive binding sites for epigenetic activators (MED1 and p300) ^[38,39]^, signal effectors (β-catenin, TCF-1, and SMAD4) ^[40–42]^, and master regulators (T and SOX2) (Figure 1e; Figure S1f, Supporting Information) ^[43,44]^. Published gene run-on sequencing (GRO-seq) ^[45]^ using *in vitro* PS counterpart also demonstrated the existence of nascent transcripts from the DRE-1 locus (Figure 1e). Given the high tissue-specificity of DRE-1 in the posterior tissue, we named the element as posterior enhancer (p-Enh) in the following studies. Consistent with the GRO-seq and eRNA quantification results (Figure 1c, e), RNA *in situ* visualization using RNAscope revealed that only sense transcripts (oriented identically to the nearby gene) could be specifically detected in PS and adjacent epiblast region of the E7.5 embryos (Figure 1f; Figure S1g, Supporting Information). There was no eRNA signals could be detected in ExE cells, PS cells of earlier embryonic stages (E6.5 and E7.0), or in the tailbud of the organogenetic embryos (E9.5) (Figure 1f; Figure S1h, i, Supporting Information). Additionally, we verified the reliability of the eRNA detection method. Despite observing robust expression of the *Cdx2* in the ExE region of the E6.5 embryo, we found no signals of eRNA (Figure S1j, Supporting Information). This result eliminated the possibility of false inferences due to limited unspliced intronic fragments of newly transcribed *Cdx2*.

Taken together, we identified a DRE, p-Enh, which is pre-marked by H3K27ac specifically in posterior epiblast at E7.5, possessing the posterior tissue-specific enhancer activity from late gastrulation stage to organogenesis stage. Notably, p-Enh shows transient sense eRNA transcription with strict spatial-temporal specificity, occurring only in E7.5 posterior epiblast.

### 2.2. Genetic removal of p-Enh causes severe posterior tissue developmental defects and embryonic lethality at the organogenetic stage

To investigate the potential functional role of p-Enh during mouse embryo development, we genetically knockout the p-Enh (∼1.3 kb) in mouse embryo using CRISPR/Cas9 system (Figure S2a, Supporting Information). No live mouse individual with homozygous p-Enh removal (p-Enh-KO) could be acquired (Figure S2b, Supporting Information). Further analyses identified that most p-Enh-KO embryos are deceased around the E10.5-E11.5 stage with noticeable posterior tailbud developmental abnormalities and hindlimb absence (Figure 2a; Figure S2c, d, Supporting Information). TUNEL assay further confirmed that the apoptotic signals become prominent in the E10.5 p-Enh-KO embryos, predominantly enriched at the tailbud and the brain regions (Figure S2e, Supporting Information).

**Figure 2.**
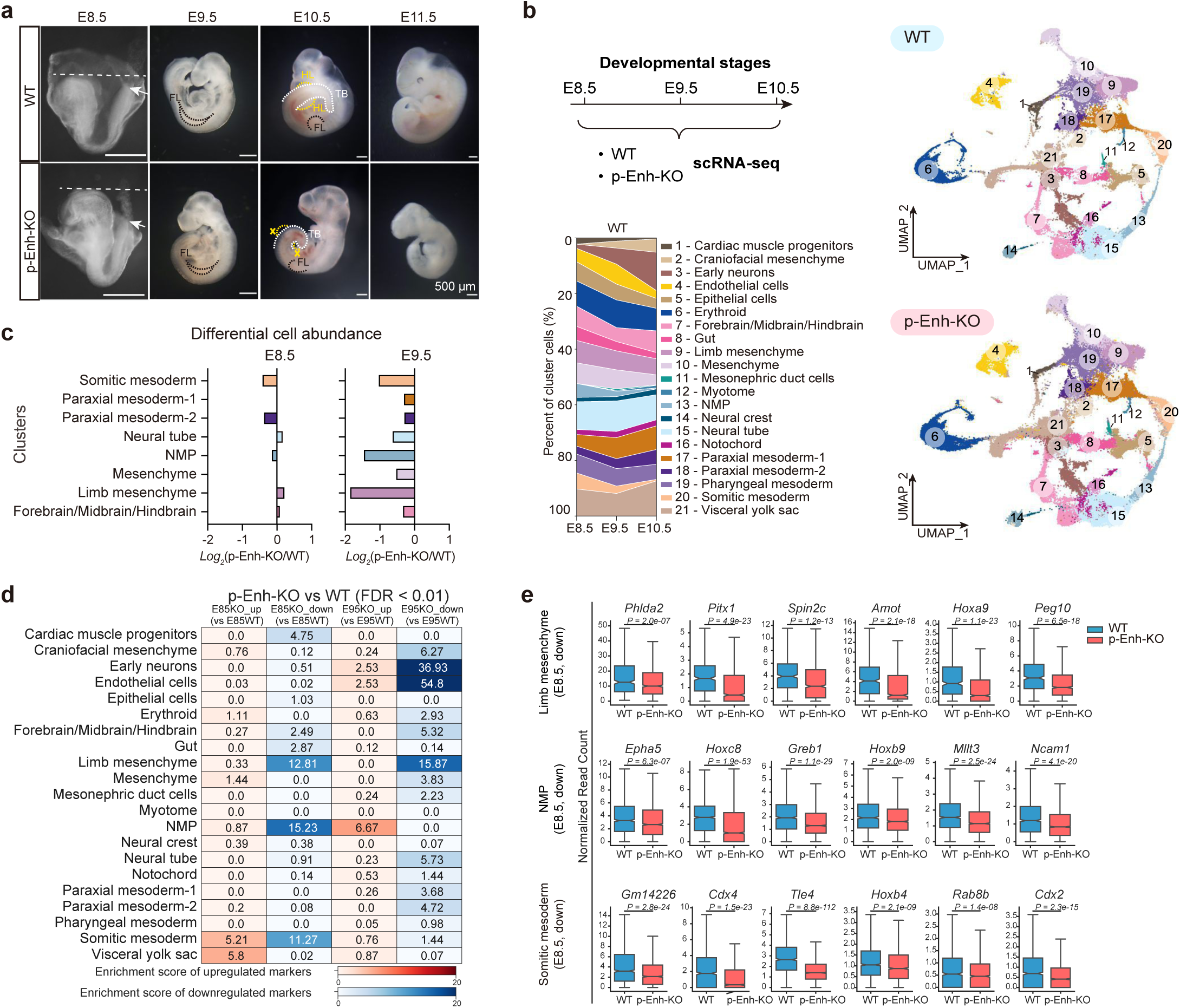
Genetic removal of p-Enh causes severe posterior tissue developmental defects and embryonic lethality at the organogenetic stage. **(a)** Bright field images of WT and p-Enh-KO mouse embryos at different development stages (E8.5, E9.5, E10.5, and E11.5). The white dotted lines representing the top baseline of the head. Arrowheads indicating the tailbud region. The tailbud and limb were outlined with dotted lines, with tailbud in white, forelimb in black, and hindlimb in yellow. FL: forelimb; HL: hindlimb; TB: tailbud. Scale bar: 500 μm. **(b)** Top left: the schematic diagram describing the sampling strategy of single-cell RNA-seq. Right: UMAP plots of single-cell RNA-seq data from WT and p-Enh-KO embryo from E8.5 to E10.5, color-coded by cluster identities. Bottom left: the percent of cluster cells in WT embryos displaying progressive cell-type complexity changes. **(c)** Differential abundance of cell clusters with pronounced changes in p-Enh-KO compared with WT based on log_2_(fold changes). **(d)** Enrichment score illustrating expression of top-100 marker genes in each cell clusters (Fisher’s exact test, FDR<0.01). The enrichment score of upregulated genes were labeled with red color, and the downregulated genes were in blue. **(e)** Boxplot illustrating normalized read count of marker genes with significant expression changes between WT and p-Enh-KO embryos at E8.5 in indicated clusters. p-value was indicated in each gene.

To systematically characterize the molecular phenotypes of p-Enh-KO embryos, we performed single-cell transcriptomic analyses in p-Enh-KO and wildtype counterparts from the same litter covering successive developmental stages from E8.5 to E10.5 (Figure 2b). In total, 180,930 cells across three sequential stages were sequenced with a median coverage of 23,783 unique molecular identifiers (UMIs) per cell. 127,069 cells were retained for further analysis after rigorous quality filters (Figure S2f, Supporting Information; Experimental Section). By referencing the published embryo atlas dataset ^[46,47]^, 21 cell clusters of the WT embryos with precise annotation were identified (Figure 2b; Figure S2g, Supporting Information). Generally, our annotation unveiled diverse clusters with unique set of markers spanning all three germ layer derivatives. For example, the neural tube cells were defined by the expression of *Sox2, Pax6, and Fabp7*, while NMPs were distinguished by *Epha5*, *T, and Fgf8* (Figure S2g, Supporting Information). Additionally, the dynamics of cell type proportions across developmental stages also reflect the sequential cell fate differentiation during organogenesis (Figure 2b). Thus, the single-cell transcriptome profile of early organogenesis ranging from E8.5 to E10.5 faithfully captures the *in vivo* developmental events.

Next, we projected the single-cell transcriptome of p-Enh-KO embryos onto the WT reference, and found that the major cell type compositions between the WT and p-Enh-KO embryos were comparable (Figure 2b). The prominent apoptotic signals identified in E10.5 p-Enh-KO embryos were further confirmed by the assessment of apoptosis and hypoxia activity hallmarks expression (Figure S3a, Supporting Information). To mitigate the potential bias caused by apoptotic cells, we focused our analyses on the E8.5 and E9.5 embryos in the following sections. Differential cellular abundance analysis highlighted a substantial decrease in posterior derived tissues of p-Enh-KO embryos, such as NMP, somitic mesoderm, and limb mesenchyme (Figure 2c; Figure S3b, Supporting Information). Following this, we calculated the enrichment scores of marker gene sets within each cluster. Remarkably, while the most significant cell density changes were observed at E9.5 (Figure 2c), the marker gene expression patterns, especially for NMP and somitic cells, showed severe abnormalities as early as E8.5 in p-Enh-KO embryos (Figure 2d, e; Figure S3c, Supporting Information). The alterations of gene expression prior to those of cellular phenotypes indicates that p-Enh may function at an earlier stage when the posterior tissues were differentiated from the PS region in the gastrula.

### 2.3. Developmental anomalies in both A-P axis patterning and limb development in p-Enh-KO embryos

Given the phenotypes of p-Enh-KO were mainly associated with the posterior tissue development, we focused on cell types comprising the embryonic A-P axis from the single-cell reference atlas, encompassing forebrain/midbrain/hindbrain, neural tube, NMP, somitic mesoderm, and paraxial mesoderm. Sub-clustering of these single cells by UMAP further divided the cells into 7 clusters (Figure 3a). RNA velocity and pseudo-time transcriptional ordering analyses showed a clear bi-directional trajectory originated from NMP towards the neural and mesodermal lineages, respectively (Figure 3a). This result aligns with the model that two distinct developmental trajectories exist in neural tube cells: one originating from the anterior brain cells and the other from the bipotential NMP in the posterior region ^[27,28]^. The developmental signal gradients (RA, Wnt, and FGF signaling) and A-P axis-related marker genes alignment ^[48,49]^ signify the robust correlation between the plotted single-cell transcriptome and the A-P axis patterning process *in vivo* (Figure S4a-d, Supporting Information), suggesting that the reconstructed UMAP plot could effectively represent cellular distribution along the *in vivo* A-P axis, and highlight NMP as the origin for posterior neural tube cells (Figure 3a).

**Figure 3.**
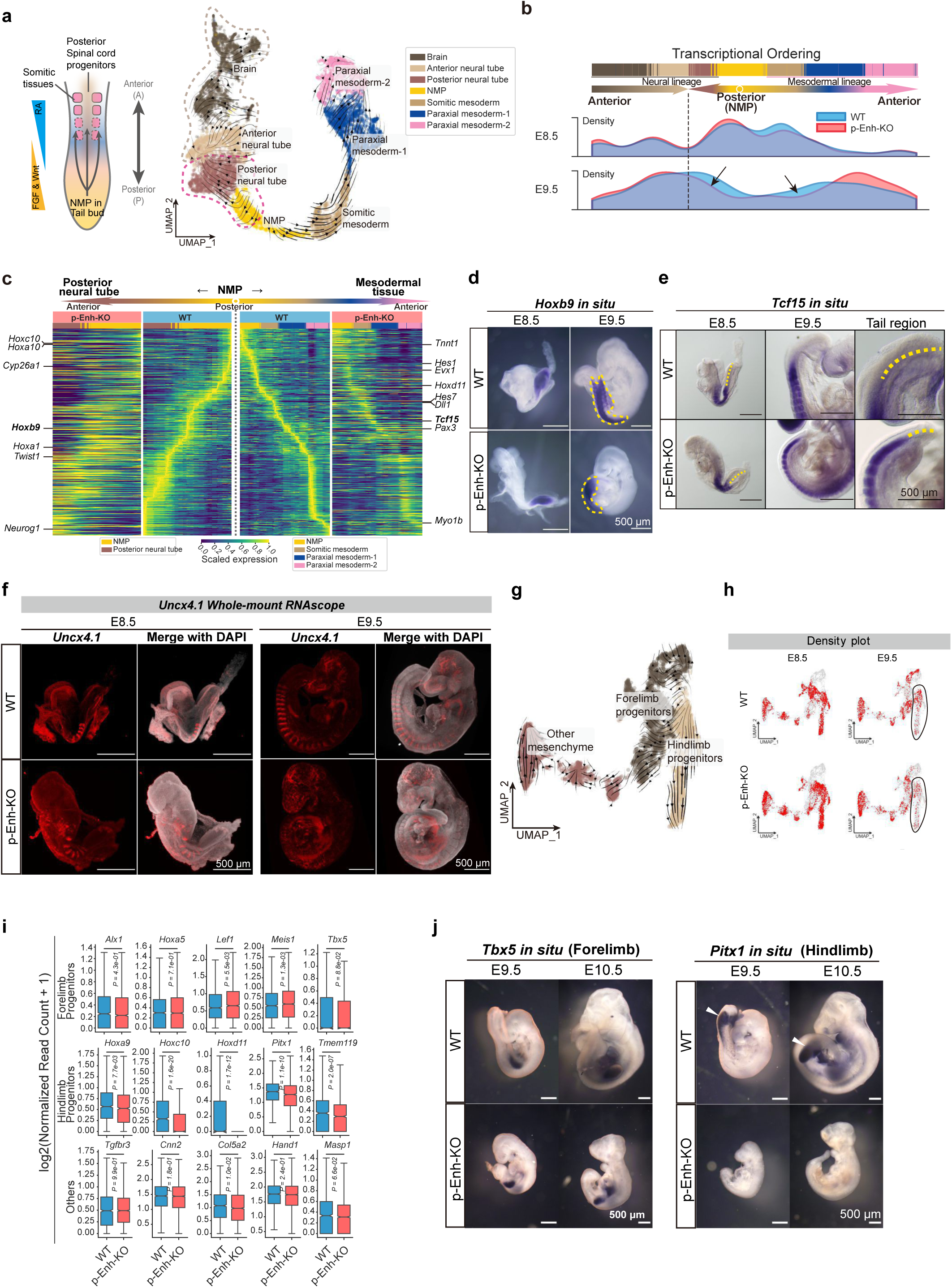
Developmental anomalies in both A-P axis patterning and limb development in p-Enh-KO embryos. **(a)** Left: cartoon diagram demonstrating the structure and cell types in tailbud region. RA, Wnt, and FGF signaling showed gradient activities along the A-P axis. Right: UMAP visualizing the subclusters and RNA velocity projection field. The dashed lines delineated diverse developmental trajectories within the neural tube. **(b)** Cell density distribution of the A-P axis related cells along one-dimensional transcriptional axis. The dashed lines indicating boundary between anterior-derived and NMP-derived neural tube cells. The black arrows indicating noticeable cell density decrease in p-Enh-KO embryos compared with WT. **(c)** Heatmap showing scaled expression of trajectory-based expressed genes based on NMP-derived tissues (NMP, posterior neural tube, somitic mesoderm, and paraxial mesoderm-1, -2). Examples of key TFs were labeled. **(d)** *In situ* hybridization results showing the expression patterns of *Hoxb9* in WT and p-Enh-KO embryos at different developmental stages. Dashed lines outlining the *Hoxb9*-positive regions in posterior neural tube and somites. Similar results were obtained from embryo replicates. Scale bar: 500 μm.3 3 **(e)** Expression patterns of *Tcf15* in WT and p-Enh-KO embryos at different developmental stages indicating shrinkage of the undifferentiated progenitor of PSM. Yellow dashed lines indicating *Tcf15*-negative tailbud region. Similar results were obtained from embryo replicates. Scale bar: 500 μm. **(f)** Lightsheet imaging of whole-mount RNAscope targeting *Uncx4.1* in WT and p-Enh-KO embryo samples at E8.5 and E9.5. The clear strip-like pattern of *Uncx4.1* in WT embryos disturbed in E9.5 p-Enh-KO embryos. Scale bar: 500 μm. **(g)** UMAP visualizing the projection field of limb related cell clusters with RNA velocity. Colors indicated different cell types. **(h)** Density plots illustrating the distribution of limb related cells in WT and p-Enh-KO embryos at different developmental stages. **(i)** Boxplot showing the expression (log_2_(Normalized Read Count +1)) of limb marker genes in WT and p-Enh-KO samples. p-value was labeled on each boxplot. **(j)** *In situ* hybridization of forelimb marker *Tbx5* and hindlimb marker *Pitx1* in WT and p-Enh-KO embryos. Scale bar: 500 μm.

Subsequently, we compared the cellular distribution between wildtype and p-Enh-KO embryos along with the reconstructed digital A-P map. By linearly plotting the single cells along the reconstructed A-P axis, the p-Enh-KO embryo exhibited a noticeable reduction in cell density in NMP and their immediate developmental progenies (Figure 3b). Trajectory-based gene expression analyses revealed significant disruptions in the developmental programs during both posterior neural tube and mesodermal lineage development in p-Enh-KO embryos (Figure 3c). Experimental validation showed that the expression territory and abundance of *Hoxb9,* a marker for posterior neural tube and mesodermal cells with the M-shaped distribution, was restricted in the tailbud tip of the p-Enh-KO embryos (Figure 3d). Additionally, *Tcf15*, marker for anterior presomitic mesoderm (PSM) and somite ^[50]^, exhibited a substantial decrease and shrinkage in the undifferentiated progenitor pool for PSM (*Tcf15*-negative region) in E9.5 p-Enh-KO embryos (Figure 3e). Besides, the highly organized stripe-like pattern of *Uncx4.1* in WT embryos ^[51]^ was severely disrupted in p-Enh-KO embryos, especially for embryos at E9.5 (Figure 3f). Interestingly, the anterior somite structure appears to be normal in the E8.5 p-Enh-KO embryos (Figure 3f). Given the different trajectory routes for the early-anterior (early PS-derived) and late-posterior somitogenesis (NMP-derived) during mouse embryonic development ^[24]^, the disrupted *Uncx4.1* distribution indicated that p-Enh may specifically regulate the NMP-derived late-posterior somitogenesis process in the mouse embryo.

Additionally, we also observed a notable decrease in the posterior-derived limb mesenchyme cell population in E9.5 p-Enh-KO embryos (Figure 2c). To delve into these abnormalities, we specifically analyzed cells associated with limb mesenchyme and further categorized them into three sub-clusters representing forelimb progenitors, hindlimb progenitors, and other mesenchyme based on marker gene expression. RNA velocity analysis confirmed that both forelimb and hindlimb progenitors seem to share one common developmental origin (Figure 3g). Cell density analyses reveal comparable patterns between WT and p-Enh-KO embryos at E8.5, whereas there is a pronounced reduction in hindlimb progenitors in E9.5 p-Enh-KO embryos (Figure 3h). Molecular analyses identified that genes related to hindlimb development, such as *Hoxc10*, *Hoxd11*, *Pitx1*, and *Tmem119*, were significantly down-regulated in p-Enh-KO embryos, while the markers of forelimb showed limited expression alterations (Figure 3i). Further experimental validation of the typical hindlimb marker *Pitx1* confirmed the absence of hindlimb structure in the p-Enh-KO embryos with the normal distribution of forelimb cells as marked by *Tbx5* (Figure 3j).

Together, these results indicate that the deletion of p-Enh had a profound impact on the developmental programs of multiple organs, including retardation in posterior tissue growth and eventual embryonic lethality. Single-cell transcriptomics analysis reveals that p-Enh can regulate the gene expression of NMP-derived clusters as early as E8.5, p-Enh-KO results in a pronounced decrease in cell density in posterior cell clusters at E9.5, and ultimately contributing to the lethality phenotype at E10.5.

### 2.4. Two distinct mechanisms with differential CDX2 dependencies underlying p-Enh function

A substantial body of work on various enhancers have proposed mechanisms of enhancer function by regulating adjacent genes ^[52–56]^. Considering that p-Enh is located in the first intron of *Cdx2* and the crucial role of *Cdx2* in body axis development ^[31,57–59]^, we examined whether the expression of *Cdx2* was affected in p-Enh-KO embryos. There was no change in *Cdx2* expression in the extraembryonic cells, with partial down-regulation in the posterior regions at E7.5 (Figure 4a). Consistently, single-cell transcriptome data also showed reduced *Cdx2* expression in posterior tissues such as NMP and somitic mesoderm (Figure S5a, Supporting Information). To probe the downstream molecular mechanisms for p-Enh function and dissect the potential relevance between p-Enh and its neighboring coding gene-*Cdx2*, we prepared the p-Enh-KO and Cdx2-KO mouse embryonic stem cell lines with CRISPR/Cas9 system, respectively (Figure S5b, Supporting Information) ^[60]^. It is noteworthy that the Cdx2-KO cell line features a specific 35 bp deletion in the first exon, causing a frameshift mutation and eliminating functional CDX2 protein without interrupting the p-Enh element genetically (Figure S5c, d, Supporting Information). Generally, the morphologies of stem cell clones and distributions of pluripotent markers NANOG and OCT4 were comparable among WT, Cdx2-KO, and p-Enh-KO ESCs (Figure S5e, Supporting Information), indicating that the removal of CDX2 protein or p-Enh element did not affect stem cell pluripotency. Next, we applied the WT, Cdx2-KO and p-Enh-KO cell lines to the newly developed *in vitro* gastruloids ^[61]^ and Trunk-like structures (TLSs) ^[62]^ differentiation system (Figure 4b), which can be used to model the anterior-posterior axis formation and later somitogenesis in the embryo. We found that the WT embryoids could generate tailbud-like structures that resemble the caudal region of embryos in both gastruloids and TLSs system (Figure 4c; Figure S5f, Supporting Information). However, the elongation of tailbud regions in both the gastruloids and TLSs for Cdx2-KO and the p-Enh-KO cells was largely compromised in comparison to the WT group. Moreover, in contrast to the WT and Cdx2-KO gastruloids and TLSs, the embryoids derived from p-Enh-KO ESCs remain as large spherical aggregates with no morphological evidence of A-P axis elongation (Figure 4c; Figure S5f, Supporting Information). Geometric analysis of the acquired gastruloids and TLSs further confirmed the consistent and significant defects (p<0.0001) in A-P axis organization and elongation in both Cdx2-KO and p-Enh-KO embryoids. Moreover, the phenotypic abnormalities were more pronounced in p-Enh-KO embryoids compared to Cdx2-KO counterparts (Figure 4d; Figure S5g, Supporting Information).

**Figure 4.**
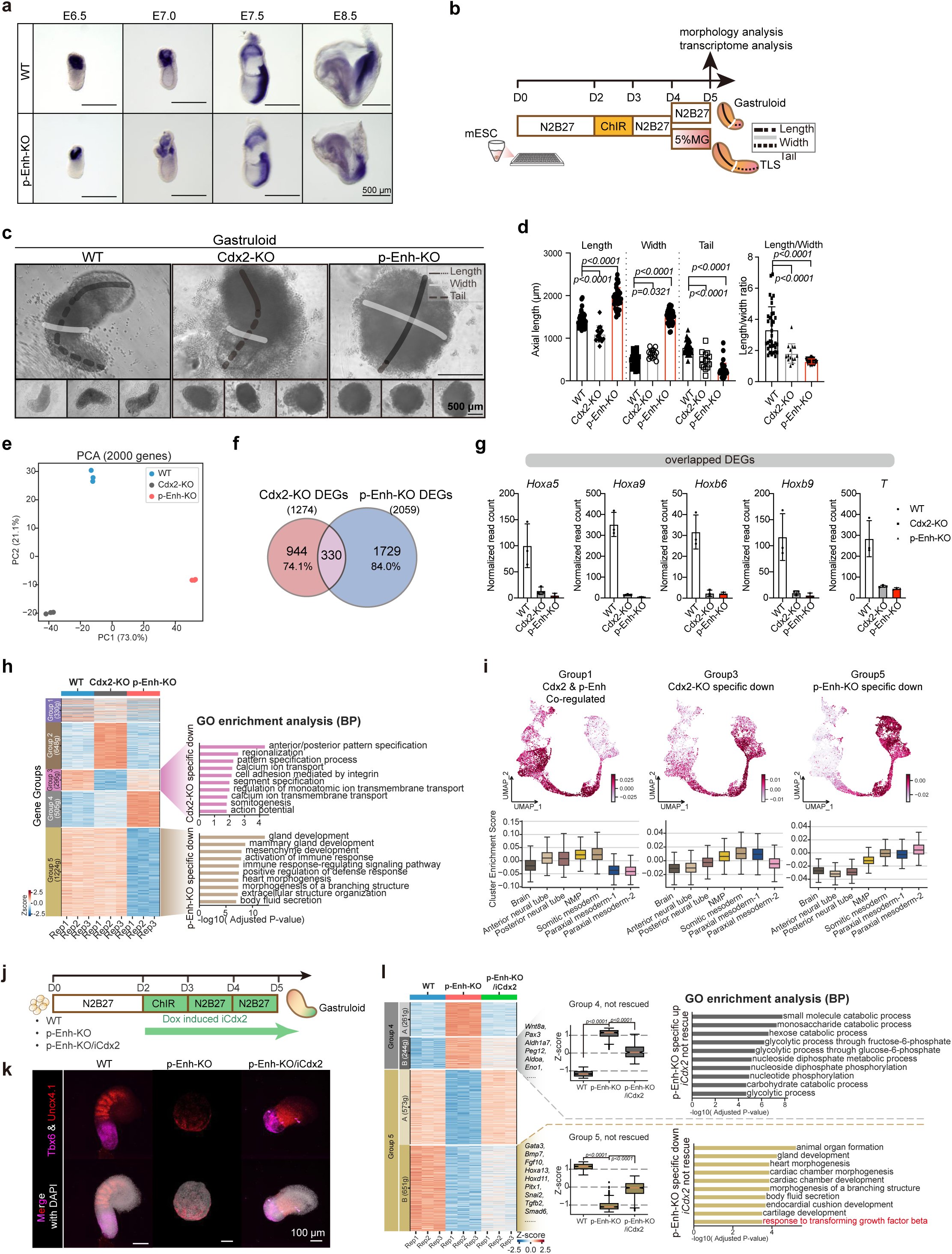
Two distinct mechanisms with differential CDX2 dependencies underlying p-Enh function. **(a)** The dynamic expression pattern of *Cdx2* in WT and p-Enh-KO embryos at different development stages. Similar results were obtained from embryo replicates. Scale bar: 500 μm. **(b)** Schematic overview of gastruloid and TLS differentiation system. The length, width, and tail were indicated, which were measured in Figure. 4c and Extended Data Figure 5f for the statistical analysis. **(c)** Representative bright-field images of gastruloids at D5 from WT, Cdx2-KO, and p-Enh-KO groups. The length, width, and tail were labeled and measured for morphological assessment. Scale bar: 500 μm. **(d)** The geometric summary of different parameters of each genotype. WT (n=35); Cdx2-KO (n=26); p-Enh-KO (n=38). The p-value were calculated based on one-way ANOVA. **(e)** Principal component analysis (PCA) of transcriptome of WT, Cdx2-KO and p-Enh-KO gastruloid. Three biological replicates were used. **(f)** Venn plot showing a modest percentage of shared genes between Cdx2-KO affecting genes and p-Enh-KO affecting genes (adjusted p-value<0.01; fold change >2). **(g)** Examples of Cdx2-KO and p-Enh-KO overlapped DEGs. **(h)** Heatmap showing differential expressed genes in WT, Cdx2-KO, and p-Enh-KO groups. Significant terms of Gene Ontology (GO) in Group3 and Group5 were shown. **(i)** Transcriptome projection of genes regulated by *Cdx2* and/or p-Enh onto *in vivo* single cell dataset. The cluster enrichment scores were listed below. **(j)** The schematic diagram illustrating the experimental design of Dox-induced *iCdx2* in gastruloid system. The time window for adding Dox is highlighted in green. **(k)** RNAscope results targeting *Tbx6* and *Uncx4.1* in gastruloids with different genotypes. Strip-like pattern of *Uncx4.1* only appeared in WT gastruloids and could not be rescued by *iCdx2* overexpression. Scale bar: 100 μm. **(l)** Heatmap showing differential expressed genes in WT, p-Enh-KO and p-Enh-KO/iCdx2 groups. Gene examples are listed. Boxplot showing the average expression of genes across different genotypes. The Gene Ontology (Biological Process) of Group 4-B and Group 5-B are shown on the right.

To comprehensively determine the molecular architectures of gastruloids derived from Cdx2-KO and p-Enh-KO cells, we performed transcriptomic profiling of the gastruloids with three biological replicates for each group (Figure S6a, Supporting Information). Our analysis revealed a significant decrease in *Cdx2* expression in both Cdx2-KO and p-Enh-KO samples, which was further experimentally confirmed (Figure S6b, c, Supporting Information). Principal component analyses (PCA) revealed distinct transcriptomic features for Cdx2-KO and p-Enh-KO samples (Figure 4e). Additionally, p-Enh-KO exhibited a greater number of differentially expressed genes (DEGs) compared to Cdx2-KO (p-Enh-KO: 608 up and 1,451 down, vs. Cdx2-KO: 819 up and 455 down) (Figure S6d, Supporting Information). Remarkably, only a small portion of DEGs were overlapped between Cdx2-KO and p-Enh-KO group (330/1,274, 25.9% of Cdx2-KO DEGs; 330/2,059, 16% of p-Enh-KO DEGs) (Figure 4f), suggesting both shared and distinct regulatory roles of *Cdx2* and p-Enh during A-P axis formation and elongation in embryoids. Despite constituting a small fraction, the shared target genes among the overlapping DEGs, including regulators of A-P axis patterning such as *Hox* genes and *T* ^[31,63,64]^, exhibited substantial downregulation in both Cdx2-KO and p-Enh-KO gastruloids (Figure 4g). Furthermore, the majority of DEGs (84%) that do not overlap with Cdx2-KO are of particular importance to our investigation. Next, we categorized these DEGs (Figure 4f) into 5 clusters based on gene expression patterns, including the shared DEGs (Group 1), Cdx2-KO specific up (Group 2), Cdx2-KO specific down (Group 3), p-Enh-KO specific up (Group 4), and p-Enh-KO specific down (Group 5) (Figure 4h). Gene ontology (GO) analysis revealed that genes in Group 3 were enriched in the biological process related to anterior/posterior patterning and somitogenesis process, consistent with the well-known functions of CDX2 protein ^[57]^. As for p-Enh-KO specific down (Group 5) genes, the pathways related to a broad spectrum of mesodermal development, such as gland development, mesenchyme development, and heart morphogenesis, were enriched (Figure 4h; Figure S6e, Supporting Information). Following this, we conducted an enrichment analysis of the DEGs of Group 1, 3, 5 with the constructed *in vivo* single-cell A-P atlas (Figure 3a), and found that Group 1 and Group 3 exhibited a clear posteriorized enrichment, displaying high enrichment scores with NMP, somitic mesoderm, and posterior neural tube clusters. However, genes in the p-Enh-KO specific down group (Group 5) showed a strong correlation with NMP and the mesodermal clusters (Figure 4i), indicating p-Enh may participate into posterior mesoderm development through a CDX2-independent mechanism.

To further validate this notion, we established inducible *Cdx2* overexpression cell line on the background of p-Enh-KO (p-Enh-KO/iCdx2) and subject this cell line into gastruloid differentiation system (Figure 4j; Figure S6f, Supporting Information). We observed that the expression of the somitic mesoderm marker *Tbx6*, a downstream target of *Cdx2*, was reactivated in the p-Enh-KO/iCdx2 gastruloids. However, the A-P elongated morphology as well as the stripe-like structure marked by *Uncx4.1* transcripts remained compromised in p-Enh-KO/iCdx2 (Figure 4k). Transcriptome analyses showed that more than 50% of p-Enh-KO specific down genes (651/1,224, 53% of Group 5) failed to be fully rescued by the inducible *Cdx2* overexpression in comparison with the WT counterpart (Figure 4l; Figure S6g, Supporting Information), which enriched organ formation-related genes. In particular, genes involved in the response to the transforming growth factor beta (TGF-β) pathway remains abnormal even with the restoration of CDX2 (Figure 4l). Regulon analyses summarizing the upstream regulators for the un-rescuable genes revealed that major effectors involved in TGF-β signaling, such as Smad1 and Smad9, were specifically enriched (Figure S6h, Supporting Information).

In summary, *in vitro* embryoid data showed that p-Enh can regulate posterior tissue development partially through the neighboring gene *Cdx2*. Moreover, these results also suggest that a previously unrecognized CDX2-independent mechanism may contribute to p-Enh function, possibly involving cellular responsiveness to TGF-β signaling pathway.

### 2.5. Genome-wide epigenomic remodeling suggests a potential functional relevance between p-Enh and TGF-β signaling

To systematically determine the downstream molecular cascades of p-Enh, we profiled the H3K27ac modification enrichment from both WT and p-Enh-KO NMP cells acquired from directed NMP differentiation system (Figure 5a) ^[65]^. The comparative analysis of chromatin activities marked by H3K27ac revealed the occurrence of pervasive epigenomic remodeling in p-Enh-KO NMP cells, especially at distal intergenic chromatin regions (Figure 5b; Figure S7a-c, Supporting Information). GREAT analyses identified that peaks with weakened enrichment of H3K27ac in p-Enh-KO cells are strongly enriched with SMAD binding capabilities (Figure 5b), which are crucial to mediate the responsiveness to TGF-β signaling ^[8]^. Following this, we also incorporated published dataset of SMADs ChIP-seq data ^[66,67]^, as well as CDX2 ChIP-seq ^[31]^ data from *in vitro* sample counterpart, and checked the distribution of these signaling effectors around peak sets with changed H3K27ac level in p-Enh-KO cells (Figure 5c). We observed enrichment of TGF-β signal effectors (SMADs) binding with these peaks, but with minimal CDX2 binding (Figure 5c). This result was further supported by the examination of developmental gene loci related to A-P axis patterning such as *Tbx3* and *Tcf15*, as well as genes involved in developmental signaling pathway, like *Tgfbr2* (Figure S7d, Supporting Information). These results, together with previous transcriptome data (Figure 4f, l), indicate that the CDX2-independent mechanism underlying p-Enh function may participate in coordinating with TGF-β signaling.

**Figure 5.**
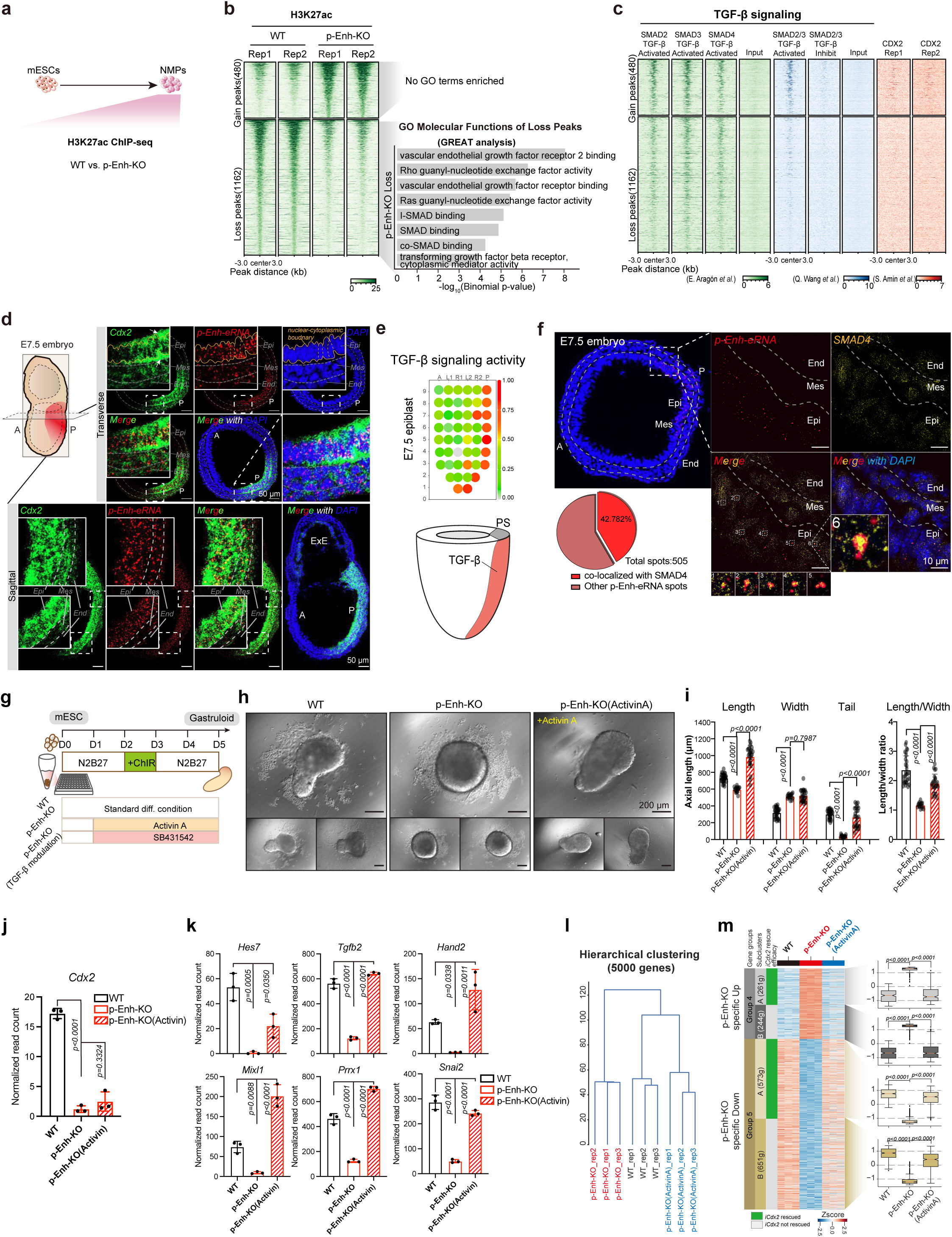
Functional interplay between p-Enh and the TGF-β signaling pathway. **(a)** Schematic of 2D NMP differentiation system. H3K27ac ChIP-seq data were employed to assess the chromatin activity of WT and p-Enh-KO NMPs. **(b)** Heatmap showing differential H3K27ac peaks in WT and p-Enh-KO NMPs. GO terms of top-ranked GREAT analysis from differential H3K27ac peaks were shown. No GO terms enriched in p-Enh-KO gain group under default parameters. **(c)** Heatmaps illustrating chromatin binding patterns of multiple TGF-β signaling downstream effectors from two independent studies. SMAD2, SMAD3, SMAD4 in TGF-β-activated condition ^[66]^, SMAD2/3 in TGF-β-activated condition ^[67]^. Control: CDX2 ChIP-seq ^[31]^. **(d)** Co-staining of *Cdx2* mRNA and p-Enh-eRNA using RNAscope in E7.5 mouse embryos. Both transverse and sagittal sections were shown. The labeling scheme was consistent with Figure 1f, and the nuclear-cytoplasmic boundary was outlined in a vibrant yellow color. Scale bar: 50 μm. **(e)** TGF-β signaling activity determined by gene set (GESA: MM9770) during gastrulation. The schematic diagram was adapted and modified from the spatial transcriptome of gastrulation ^[9]^. **(f)** RNAscope targeting p-Enh-eRNA and co-staining with SMAD4 protein in E7.5 mouse embryo sections. Zoomed-in view showing the detailed signals at posterior region. The labeling scheme was consistent with Figure 1f. The co-localization spots of p-Enh-eRNA with SMAD4 proteins were further magnified for the clear presentation. The co-localization analysis was performed using ImageJ plugin ComDet 0.5.5. Scale bar: 10 μm. **(g)** Diagram showing modulation of TGF-β in gastruloid differentiation system. Activin A (20 ng/mL) or SB421542 (10 μM) were added into differentiation system from the end of D1. **(h)** Bright-field images of D5 gastruloids in different conditions. Scale bar: 200 μm. **(i)** Geometric features (length, width, tail length, and length/width ratio) of D5 gastruloids in Figure 5h. The p-value were calculated based on one-way ANOVA. **(j-k)** Expression pattern of *Cdx2* and example genes in WT, p-Enh-KO and p-Enh-KO (Activin) gastruloids. The p-value were calculated based on one-way ANOVA. **(l)** Hierarchical clustering of transcriptome from WT, p-Enh-KO and p-Enh-KO (ActivinA) gastruloids. **(m)** Heatmap illustrating expression pattern of p-Enh specific regulated genes (defined in Figure 4h, l) in different groups. Subclusters A and B were delineated according to the rescue efficacy of *iCdx2* overexpression. Boxplot showing the average expression of genes in each subclusters.

Recent studies have found that a broad spectrum of TFs bind with RNA molecules, highlighting the conserved and essential nature of interactions between TFs and RNA in vertebrate development ^[68]^. We found that p-Enh can transcribe into RNAs, which are largely distributed in the posterior regions at the late-gastrulation stage (E7.5) (Figure 1f). Co-detection of p-Enh transcripts and *Cdx2* mRNA in the E7.5 mouse gastrula revealed that both transcripts were specifically enriched in the posterior region (Figure 5d). Moreover, sub-cellular distribution analyses showed that the p-Enh-eRNAs were prominently enriched in the nuclei, forming puncta-like structure at multiple loci within a single nucleus. In contrast, the transcripts of the neighboring coding-gene, *Cdx2,* were universally located in the cytoplasm (Figure 5d). Meanwhile, the signaling activity of TGF-β was also enriched in the posterior embryonic region at E7.5, represented by the spatial gene expression enrichment of typical TGF-β signaling target genes (Figure 5e). Co-staining of p-Enh-eRNA and SMAD4 protein in the E7.5 mouse embryo revealed that more than 40% nuclei-enriched p-Enh-eRNA puncta were specifically co-localized with SMAD4 protein (Figure 5f). The spatial co-occurrence of p-Enh transcripts and TGF-β signaling activity hints an intriguing involvement role of p-Enh-eRNA in TGF-β signaling transduction.

To further validate the involvement of TGF-β in p-Enh’s functions, we activated TGF-β signaling using Activin A in gastruloid differentiation system (Figure 5g; Figure S8a, Supporting Information). Interestingly, activation of TGF-β signaling in p-Enh-KO cells markedly ameliorated the differentiation defects (Figure 5h, i). Transcriptome analysis following TGF-β activation in gastruloids indicated that the rescue effect was not mediated through the neighboring gene *Cdx2*, which remained low level even after Activin A treatment (Figure 5j). This supports the notion that the differentiation defects could be attributed to the interactions between p-Enh and TGF-β signaling in a *Cdx2*-independent way, as suggested in Figure 4l. In contrast to the expression pattern of *Cdx2*, a list of genes crucial for A-P axis patterning and mesoderm development, including *Hes7*, *Tgfb2* and *Mixl1*, exhibited marked upregulation upon TGF-β activation. This underscores the ability of TGF-β to effectively ameliorate the phenotype associated with p-Enh-KO (Figure 5k). Hierarchical clustering result also revealed that Activin A treatment effectively rescued genes expression abnormalities caused by p-Enh depletion (Figure 5l). Specifically, p-Enh regulated genes that failed to be rescued by *Cdx2* re-expression (subcluster B of Group 4 and Group 5 in Figure 4l) were largely restored with Activin A treatment (Figure 5m). Conversely, inhibition of TGF-β signaling with SB431542 induced extensive cell death, indicating the essential roles of TGF-β signaling in A-P axis patterning (Figure S8b, Supporting Information). Together, these findings suggest that the crosstalk between p-Enh and the TGF-β signaling pathway holds substantial biological importance for the development of posterior tissues.

## 3. Discussion

In this study, we have identified a pre-marked regulatory element, p-Enh, exhibiting highly restricted spatial-temporal activity and significant biological importance during mouse early embryonic development. The pivotal role of p-Enh in posterior tissue development is underscored by the embryonic lethality observed in p-Enh-KO embryos, accompanied by severe posterior tissue growth failure, disruption of A-P axis gene expression pattern, and dysfunctions across multiple cell types (Figure 2 and 3). Mechanism analyses through transcriptomic dissection pinpointed the co-occurrence of two distinct mechanisms underlying the p-Enh function: (i) regulating the expression level of its neighboring gene *Cdx2 in cis*, thereby affecting the downstream cascades of CDX2 (Figure 4); (ii) transient production of nuclei-distributed eRNAs, potentially involved in coordinating the nuclear responsiveness to the extrinsic developmental signals, such as TGF-β signaling pathway (Figure 5). Thus, we propose a molecular model for pre-marked developmental enhancers, wherein genetic sequence-based *in cis* local regulation and RNA transcripts-based *in trans* genome-wide regulation co-exists to facilitate tight spatiotemporal control of global molecular architecture and thus set up the molecular blueprint for future tissue development (Figure 6).

**Figure 6.**
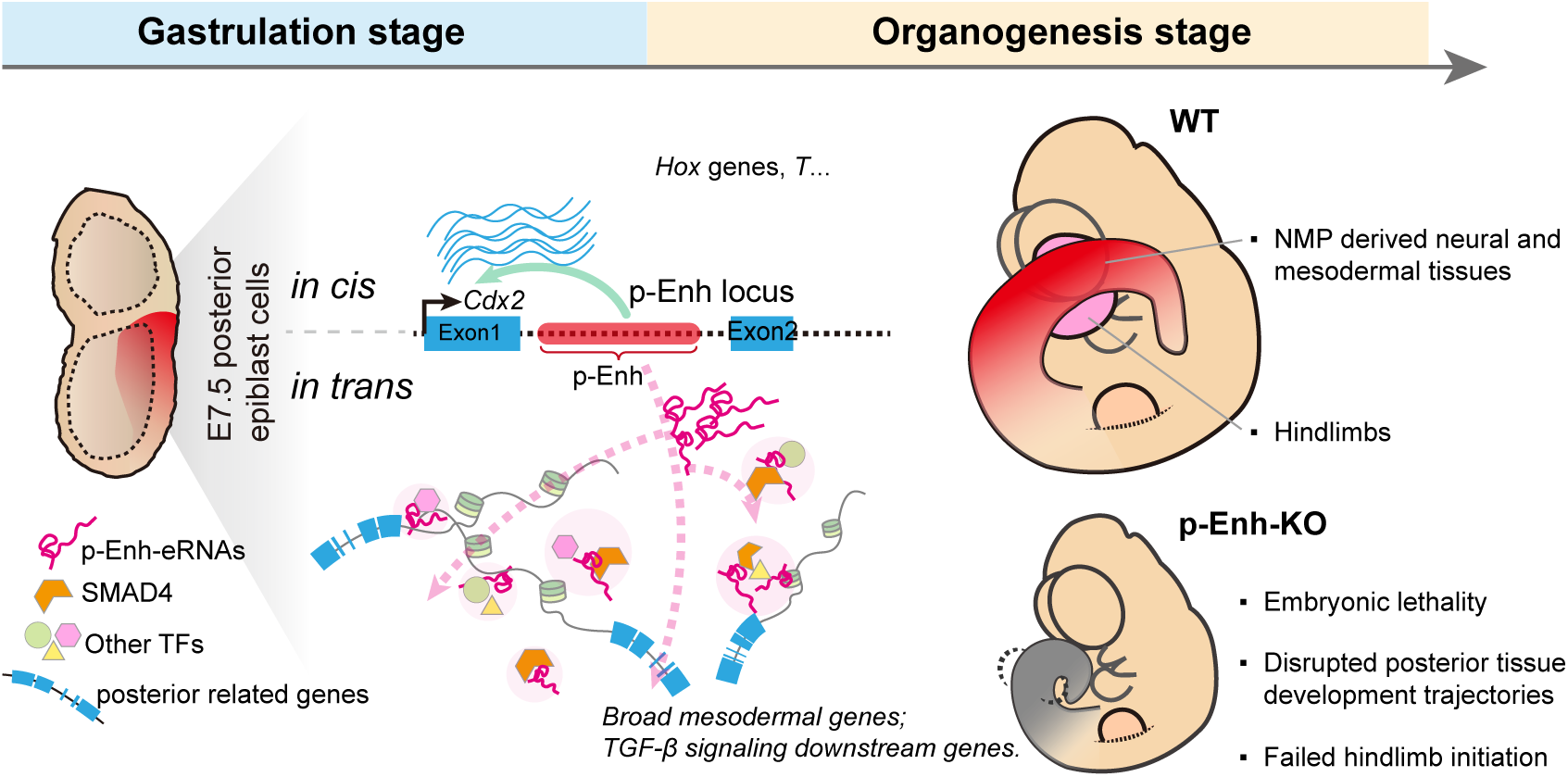
Model of the regulation of posterior developmental genes through p-Enh with distinct in cis and in trans mechanisms. The gastrula pre-marked p-Enh plays dual roles in governing posterior tissue development: (i) regulating the neighboring gene *Cdx2* (*in cis*), which in turn affects their shared downstream targets, such as *Hox* and *T* genes; (ii) modulating the global transcriptome and epigenome via transient expression of broadly nuclei-distributed eRNA (*in trans*), potentially by mediating the interplay between p-Enh-eRNAs and signaling effector such as SMAD4, which may contribute to the modulation of TGF-β signaling. p-Enh-KO embryos exhibit embryonic lethality, aberrant posterior tissue development trajectories, and failure to initiate hindlimb formation.

Recently, there has been a growing interest in dissecting the molecular hierarchy of gene regulatory networks (GRNs) that govern rapid cell fate commitment ^[9,18,69,70]^. Previous studies have primarily focused on identifying specific pioneer transcription factors capable of steering the developmental competence towards specific developmental lineages ^[71–73]^. In line with the pioneering factors, emerging evidences also indicated that the epigenetic priming or pre-mark at regulatory elements could offer a notable feature prior to cell fate decision ^[13,74]^. However, it remains unclear whether the “chromatin priming” of regulatory elements holds potential functional roles in biological process beyond merely serving as a marker. Here, we determined one pre-marked regulatory element, p-Enh, which is specifically activated in the PS and adjacent epiblast at late gastrulation stage, and sustains the epigenetic activity till late organogenesis. Phenotypic and molecular analysis demonstrated that genetic loss of p-Enh leads to dramatic developmental defects when the posterior lineage diversifies. Moreover, the precedence of transcriptomic disruptions over phenotypic anomalies strengthens the evidence for the importance of the priming effects. Thus, future evaluation of pre-marked regulatory elements and further phenotyping through incorporating multi-dimensional data may better elucidate the function of novel crucial regulatory elements.

A considerable amount of research has put forth on how enhancers exert their functions, particularly through the regulation of neighboring genes. Here, through in-depth transcriptomic dissection and genetic analyses, as reflected by the transcriptomic comparison among multiple genetic edited cell lines (WT, p-Enh-KO, Cdx2-KO, p-Enh-KO/iCdx2), we found that only 16% genes were regulated through the p-Enh-*Cdx2* axis (Figure 4f). Meanwhile, TGF-β signaling responsive genes were highly enriched in the cluster of p-Enh specifically regulated genes (Figure 4l; Figure S6h, Supporting Information). Further, through profiling H3K27ac pattern in both WT and p-Enh-KO cells, we found that the global distribution of H3K27ac was pervasively remodeled in p-Enh-KO cells (Figure 5b). Interestingly, the genomic regions with changed H3K27ac level are enriched with TGF-β signaling effectors but not CDX2 binding (Figure 5c). Thus, the regulatory mechanisms underlying p-Enh function are intricately linked with TGF-β signaling, suggesting a complex interplay rather than a straightforward linear molecular cascade through its neighboring gene *Cdx2*. p-Enh possesses both *in cis* and *in trans* regulatory mechanisms in the developmental control of posterior tissues.

The discovery of enhancer-derived transcripts largely broaden the current scope of the molecular mechanisms governing chromatin structure organization and gene expression regulation ^[75,76]^. The abundant expression and nuclear localization of p-Enh-eRNAs at multiple loci within individual nuclei in the PS and adjacent epiblast region suggest their potential roles in global GRNs instead of localized functions. Their broad distribution within nuclei may contribute to the observed pervasive chromatin activity changes in p-Enh-KO cells (Figure 5a, c). Furthermore, recent study has underscored the extensive interaction between TFs and RNA molecules ^[68]^. Drawing from the existing research and our findings, we hypothesize that the widespread eRNAs within individual nuclei might interact with signaling downstream effectors, potentially serving as “regulatory units” that modulate pivotal biological events, including the *in trans* regulatory functions of p-Enh (Figure 6). These “regulatory units” could potentially facilitate the efficient guidance of interacting TF repertoires to their target chromatin regions, bridging the enhancer element and signaling pathway targets through eRNA-TFs molecules, thereby influencing the long-range modulation of global GRNs for proper developmental programming. Future in-depth characterization of the structure of p-Enh-eRNA, its targets, and the molecular partners of this transcript will clarify the complex regulatory network and promote a comprehensive understanding of mouse embryogenesis.

In summary, to delve into the molecular mechanisms underlying epigenetic pre-marked enhancers, we systematically dissect both the *in cis* and *in trans* paths for p-Enh, proposing this as a novel strategy of developmental enhancer to coordinate multiple downstream modules genome-widely, and then establish a responsive cellular state for forthcoming lineage diversification.

## 4. Experimental Section

### Animals use, embryo staging and collection

Animal experiments were performed at the Animal Core Facility, under program project SIBCB-S308-1807-025 approved by the Institutional Animal Care and Use Committee in Center for Excellence in Molecular Cell Science, CAS, and the program project GZLAB-AUCP-2022-10-A04 approved by Institutional Animal Care and Use Committee of Guangzhou National Laboratory. Timed matings were set up between sexually mature (older than six weeks of age) of wildtype C57BL/6 or p-Enh-KO. Plugged female mice were picked after mating and marked as embryonic day 0.5 (E0.5). Female mice were sacrificed for embryos collection at specific gestational stages. Embryos were isolated from the uterus and carefully transferred into PBS in petri dishes, and extra-embryonic tissues were removed using needles under Olympus stereoscope. The developmental stage and morphology of embryos were reconfirmed according to the Theiler stage methodology. The acquired embryos were ready for further experiments including whole-mount *in situ* hybridization, iDISCO, or single cell transcriptome profiling.

### mESC maintenance and differentiation

mESC lines were maintained in standard feeder-free culture conditions which were supplemented with 3 μM CHIR99021 (Selleck Chemicals, S1263), 1 μM PD0325901(Selleck Chemicals, S1036) and 10 ng/mL mouse leukemia inhibitory factor (Millipore, ESG1107) as previously described ^[77]^. The NMP differentiation was performed as previously reported ^[65]^. In brief, 50,000 cells were seeding in N2B27 medium containing 10 ng/ml bFGF (Shanghai Pufei Biotechnology, 1106-010) in Matrigel pre-coated 35 mm dish and cultured for two days. The N2B27 medium contained a 1:1 ratio of DMEM/F12:Neurobasal medium (GIBCO) supplemented with 1xN2 (GIBCO), 1xB27 (GIBCO), 1x GlutaMax (GIBCO), 40 mg/mL BSA (Roche), penicillin/streptomycin (GIBCO) and 0.1 mM 2-mercaptoethanol. Cells were pulsed with 5 μM CHIR99021 during 24 h to 36 h. At the end of D3, the NMPs were ready for transcriptome profiling, chromatin conformation capture assays, and ChIP-seq analysis.

### Gastruloids and Trunk-like structure (TLS) generation

Gastruloids and TLSs were generated using established protocols with minor adjustments ^[62,78]^. The basic N2B27 medium contained a 1:1 ratio of DMEM/F12:Neurobasal medium (GIBCO) supplemented with 1xN2 (GIBCO), 1xB27 (GIBCO), 1x GlutaMax (GIBCO), penicillin/streptomycin (GIBCO) and 0.1 mM 2-mercaptoethanol. The initial 4 days of differentiation for both gastruloids and TLSs were consistent. In brief, 200-250 single mESC were seeded in 40 μL N2B27 medium into wells of the 96-well V bottom Ultra-low Attachment Plates (S-BIO, MS-9096VZ) and allowed to aggregated for 48 h. Subsequently, the spheres were treated with 3 μM CHIR99021 in 150 μL N2B27 medium for 24 h. After this, medium was refreshed every 24 h with the same volume of basic N2B27 medium. On the final day of differentiation, gastruloids were cultured with basic N2B27, while TLSs were embedded in 5% Growth-Factor-Reduced Matrigel (Corning, 356231) in N2B27 medium. At the end of day 5, the gastruloids and TLSs were subjected for morphology analysis, transcriptome sample collection, or immunofluorescence.

### Transgenic embryo enhancer activity screening

Transgenic embryo enhancer activity screen was performed following methods as previously reported ^[37]^. In brief, enhancer candidates with posterior epigenetic activity were selected from our published dataset and ranked according to H3K27ac increasing enrichment at posterior region from E7.0 to E7.5 mouse gastrula ^[18]^. Selected fragments were cloned from mouse genomic DNA and ligated into PB-pHsp68-LacZ plasmid using seamless cloning following the manufacturer’s instructions (Beyotime, D7010M). Then, the acquired PB-DREs-Hsp68-LacZ plasmids and PBase mRNA were microinjected into the cytoplasm of a fertilized egg with the FemtoJet Microinjection System (Eppendorf). The injected embryos were then cultured to the 2-cell stage in KSOM medium at 37 °C, 5% CO2 in standard incubator, and the 2-cell embryos were transferred to the oviduct of pseudo-pregnant ICR females and marked as 0.5 dpc. Finally, embryos were collected at appropriate stages and subjected to further genotyping and LacZ staining following the protocol previously reported ^[37]^.

### Whole-mount in situ hybridization

The mRNA expression in E6.5-E10.5 embryos was assessed by whole-mount in situ hybridization using the digoxigenin-labelled antisense RNA probes as described previously ^[79]^. Primers for amplifying probe templates are listed in Table S4, Supporting Information. In brief, embryos at different stages were collected and fixed with 4%PFA overnight and dehydrated stepwise in graded series of methanol. Embryos can be stored in methanol for a minimum overnight and up to 1 week at -20°C. Embryos were rehydrated through 75%, 50% and 25% methanol at room temperature, washed three times with DPBS and treated with 10 µg/mL proteinase K (Invitrogen, AM2548) in PTW for suitable time. After post-fixation for 30 min, approximately 500 ng of digoxigenin-labelled RNA probe was incubated with the embryo at 70°C overnight. The embryos were then washed with TBST and incubated with 1:2,000 diluted anti-digoxigenin AP antibody (Roche, 11093274910) in blocking buffer. Incubating on a rocker platform at 4°C overnight. The next day, embryos were further washed with TBST and stained with NBT/BCIP solution. Images were obtained using Olympus stereoscope.

### Haematoxylin and eosin (HE) staining

Embryo samples were fixed in 4% paraformaldehyde/PBS at 4 °C overnight and then processed for paraffin wax embedding. 7-μm thick sections were cut, dewaxed in xylene, rehydrated through an ethanol series into PBS. HE counterstaining was performed using staining kit (Beyotime, C0105) according to manufacturer’s instructions. Images were taken on Olympus VS120 microscope.

### Single cell transcriptome profiling of mouse embryos

Sexually mature p-Enh^+/-^ mice were inter-crossed and pregnant females at the indicated gestational stages (E8.5, E9.5 and E10.5) were sacrificed for embryo recovery. After dissection from the decidua, embryos with the same genotype were combined together and incubated in 200 uL TrypLE (Gibco, 12604013) at 37 ℃ with shaking (300 rpm) for about 15 minutes. The tissue pellets were manually triturated with two minutes interval. Immediately following digestion, single cells were washed twice in 1% BSA/PBS and filtered through Falcon® 40 µm Cell Strainer (Corning, 352340). Finally, cells were spun at 500 g for 5 min at 4 ℃ and resuspended in 0.04% BSA/PBS for further quality check and loading onto the 10× Chromium Controller. Processing of the samples was performed using the Chromium Single Cell 3′ library & Gel Bead Kit version 3 (10× Genomics) according to the manufacturer’s instructions. Pair-end 150 bp sequencing was performed on NovaSeq platform.

### The generation of specific fragments genomic knockout ESC line and mouse

The generation of enhancer knockout or Cdx2 knockout mESCs were performed following protocol published previously with slightly modify ^[60]^. Briefly, small guided RNAs (sgRNAs) were designed according to the instruction from Chop-chop website (http://chopchop.cbu.uib.no/). sgRNAs were annealed and cloned into linearized px330-mCherry plasmid (Addgene, Plasmid #98750). Px330-mCherry-sgRNA was transfected into E14TG2a by Lipofectamine 3000 Transfection Reagent (Invitrogen, L3000008) following the manufacturer’s instructions. The mCherry positive cells were isolated and collected by BD FACS Aria SORP for further culture for 4 to 5 days until clone formation. Single clones were picked up manually for genotyping and sanger sequencing. The potential off-target sites were tested based on the Chop-chop website’s indication. Primers used for genotyping and sgRNA sequences targeting p-Enh and *Cdx2* were listed in Table S4, Supporting Information. For the generation of p-Enh-KO mice, we employed the same sgRNAs utilized for p-Enh knockout in mESCs. Cas9 mRNA and sgRNAs with scaffolds were obtained through in vitro transcription using the MEGAscript Kit (Invitrogen). Subsequently, these Cas9 mRNAs and sgRNAs were microinjected into the cytoplasm of fertilized ova utilizing the FemtoJet Microinjection System (Eppendorf). Following microinjection, the embryos were subjected to culture under standard culture conditions (37°C, 5% CO_2_) in KSOM medium until reaching the 2-cell stage. These 2-cell embryos were then surgically transferred to the oviducts of pseudo-pregnant females. Upon reaching adulthood, the resulting offspring were subjected to genotyping analysis.

### RNA extraction and quantitative PCR (qPCR) and RNA-seq library preparation and sequencing

For the in vitro culture cells and embryo samples, total RNA was extracted using TRIzol reagent (Invitrogen, 15596018). cDNA was reverse-transcribed with FastQuant RT Super Mix (Tiangen, KR108), following the manufacturer’s instructions. Quantitative PCR analysis of mRNA levels for different markers was performed using diluted acquired cDNA with Stormstar SYBR green qPCR master mix (DBI Bioscience, DBI-143) with specific primers, and all primers used in qPCR analysis were listed in Table S4, Supporting Information. RNA-seq libraries were prepared from 300 ng pure total RNA using NEBNext Ultra II RNA Library Prep kit (NEB, E7775L). Two highly reproducible biological replicates (each with two technical replicates) were performed. Pair-end 150 bp sequencing was performed on NovaSeq platform.

### Immunofluorescence

For the cultured cells, protocol for immunofluorescence was performed as described previously ^[80]^. Briefly, cultured cells were washed twice with PBS to remove detached cells, and fixed with 4% paraformaldehyde in PBS (pH 7.3) for about 30 mins at room temperature, followed by permeabilized and blocked with blocking buffer (0.5% Triton X-100/5%BSA in PBS) for 1 h. Next, primary antibodies targeting NANOG (Abcam, ab80892. 1:200) and OCT4 (Santa Cruz, sc-5279. 1:200) were diluted in blocking buffer and incubated with samples overnight at 4 °C. Then, samples were washed three times with PBS containing 0.3% Triton-X100 (PBS-Tri), followed by incubation with secondary antibodies (1:400 diluted) in PBS-Tri for 2 hours at room temperature. After wash steps and DAPI staining, samples were imaged using Leica TCS SP8 STED confocal microscope.

For the gastruloids, immunolabeling-enabled three-dimensional imaging of solvent-cleared organs (iDISCO) was used for the whole-mount immunolabeling as previously reported ^[81]^. In brief, samples were dehydrated with methanol, bleach in chilled fresh 5% H_2_O_2_ in methanol overnight at 4 °C. After washing and rehydration, samples were incubated in blocking buffer which containing 0.2% TritonX-100/20% DMSO/0.3M glycine in PBS at 37 °C overnight. Samples were washed in PBS/0.2% Tween-20 with 10 μg/mL heparin (PTwH) for 1h twice. Next, the primary antibodies targeting CDX2 (Abcam, ab76541) were diluted in PTwH/5% DMSO/3% donkey serum (1:600) and incubated with samples at 37 °C overnight. Then, samples were washed with PTwH for 1 day and incubated with specific secondary antibodies (1:400 diluted). Finally, after washing steps and nuclear labeling, solvent-based tissue clearing was carried out which involving dichloromethane (Sigma, 270997-12X100ML) and dibenzyl ether (DBE) (Sigma, 108014-1KG). OLYMPUS IXplore SpinSR was used for the imaging.

### RNAscope

Fresh mouse embryo samples were fixed in 4% PFA/PBS at 4 °C overnight. The fixed samples were further embedded in OCT and cryo-sectioned at 20 μm thickness. The RNAscope procedure was then performed using RNAscope® Multiplex Fluorescent Reagent Kit v2 (Advanced Cell Diagnostics, 323100) according to the RNAscope® Multiplex Fluorescent Reagent Kit v2 Assay User Manual (Document Number 323100-USM). Probes targeting from Advanced Cell Diagnostics *Uncx* (Cat No. 521431), *Tbx6* (Cat No. 498251-C2), p-Enh-eRNA (Cat No. 1199081-C2), *Cdx2* (Cat No. 438921-C3) were used. For the simultaneous detection of RNA and protein in embryo samples, we followed the procedural guidelines outlined in the RNAscope® Multiplex Fluorescent v2 Assay combined with Immunofluorescence - Integrated Co-Detection Workflow (ICW) (Document Number MK 51-150). SMAD4 polyclonal antibody (Invitrogen, PA5-34806. diluted at 1:100) was used. Images were acquired using OLYMPUS IXplore SpinSR and Zeiss LSM 980 confocal microscope.

### Western blotting

The cultured cells were washed 3 times with PBS, lysed with lysis buffer and heated at 100 °C for 10 mins. Protein solution was run on SDS-PAGE and the gels were transferred to nitrocellulose membranes (GE). After blocking in 5% skimmed milk at room temperature for 1h, membranes were then incubated with primary antibodies targeting CDX2 (Abcam, ab76541) overnight at 4 °C. Next, membranes were washed 3 times with TBST and incubated with HRP-conjugated secondary antibody at room temperature for 1 h. immunoreactive bands were visualized with SperSignal West Pico Chemiluminescent Substrate (Thermo Scientific, 34579) and detected using Tanon-5500 imaging system.

### Chromatin immunoprecipitation (ChIP) and ChIP-seq libraries preparation

The protocol for chromatin immunoprecipitation (ChIP) was described previously ^[18]^. Cross-linked cells were lysed in Solution I (10 mM HEPES, pH 7.9, 0.5 % NP40, 1.5 mM MgCl2, 10 mM KCl, 0.5 mM DTT) and SDS lysis buffer (1% SDS, 10 mM EDTA, 50 mM Tris-HCl, pH 8.0), and the samples were sheared for 14 cycles (30 s on/off) in a Bioruptor Pico (Diagenode, Belgium) to achieve an average fragment size of 200-300 bp. Solubilized fragmented chromatin was immunoprecipitated with antibody against H3K27ac (Active Motif, 39133). Antibody-chromatin complexes were pulled down using Dynabeads® Protein G (Invitrogen, 10004D) on a magnetic separation rack, washed several times, and then eluted from the magnetic beads. Reverse crosslink was performed subsequently under 65 °C for at least 4 hours. ChIP-DNA was treated with RNase A and Proteinase K, and precipitated with ethanol. ChIP-DNA was finally solved in nuclease-free water and quantified using Qubit. Sequencing libraries were generated by using NEBNext Ultra DNA library preparation kit (NEB, E7370). Libraries were quality-controlled and quantified using a Qubit 2.0 Fluorometer (Life Technologies) and Agilent 2100 Bioanalyzer (Agilent Technologies). High-throughput sequencing was performed on a NovaSeq instrument.

### Quantification and statistical analysis

#### Enhancer screening based on H3K27ac ChIP-seq data

##### 1. Data processing

H3K27ac ChIP-seq data for E7.0 and E7.5 mouse embryo posterior tissues were obtained from the NCBI Gene Expression Omnibus (GEO) under accession numbers GSE98101 ^[18]^. Additionally, H3K27ac ChIP-seq data for E9.5 mouse embryo tailbud were obtained from GSE84899 ^[31]^. Processing of all ChIP-seq samples started with raw sequencing reads. Firstly, Trim Galore (version 0.4.4_dev) was used to remove adapter and low-quality sequences by trimming 3’ ends of reads ^[82]^. The resulting reads were then aligned to mouse reference genome mm10 using Bowtie (version 1.2.2) with the parameters “-- chunkmbs=512 -I=0 -X=1000 --best -m=1” ^[83]^. After removing duplicates, we performed peak calling using MACS (version 1.4.2) with the parameters “--shiftsize=100 –nomodel --keep-dup=all” ^[84]^. Subsequently, we merged all resulting peaks of samples into a consensus list of genomic regions and counted the reads within those regions using MAnorm2_utils (version 1.0.0) with the parameters “--min-peak-gap=150 --typical-bin-size=2000 --shiftsize=100 --filter=blacklist” ^[30]^. The blacklist regions of mm10 were obtained from Amemiya et al ^[85]^.

##### 2. Screening distal regulatory elements activated at E7.5 mouse embryo posterior tissues

We used MAnorm2 (version 1.2.2) to identify differential H3K27ac regions between E7.0 and E7.5 embryo posterior tissue samples with the cutoffs p-value < 0.01 and fold change > 2 ^[30]^. Distal regulatory elements activated in E7.5 mouse embryo posterior tissues were defined as the differential H3K27ac regions which both exhibited increased H3K27ac levels in E7.5 samples and located greater than 1.5kb from transcription start sites (TSSs) of genes. The top-30 most significant elements are shown in Figure 1B. The gene annotation file of mm10 was obtained from the GENCODE project ^[86]^.

### Single cell RNA sequencing data analysis

#### 1. Data processing

For each dataset, raw sequencing data was preprocessed using cellranger (10x Genomics Cell Ranger 5.0.0) with default settings, including alignment to mouse reference genome mm10, filtering, barcode counting and unique molecular identifier (UMI) counting ^[87]^. The output files (matrix.tsv.gz, barcordes.tsv.gz and features.tsv.gz) generated by cellranger were subsequently used for downstream analysis. Cell doublets were estimated by scrublet pipeline (version 0.2.3) and discarded ^[88]^. Cells with fewer than 1,000 detectable genes or with higher than 5% UMIs from mitochondrial genes were excluded. Genes detected in more than 1% total number of cells were retained for following analysis. The detailed quality control information was listed in Table S2, Supporting Information.

#### 2. Construction of mouse embryo single cell transcriptome reference map

We utilized scRNA-seq datasets from E8.5, E9.5 and E10.5 WT embryos to establish a reference map with the R package Seurat (version 4.0.1) ^[89]^. Firstly, we normalized gene expression using the NormalizeData function and identified variable genes using the FindVariableFeatures function for each dataset. Specifically, gene count matrices were normalized by the total number of UMIs per cell and then multiplied by a scale factor (10000), followed by log-transformation. After using variance-stabilizing transformation (VST), top 2000 genes sorted by variance were selected as variable genes.

Next, we integrated the datasets by identifying cell pairwise correspondences, referred to as “anchors”, between datasets based on the consistently variable genes using the FindIntegrationAnchors function ^[90]^. Batch correction was performed with the identified anchors using the IntegrateData function ^[90]^. Subsequently, principal component analysis was performed using the RunPCA function and top 30 principal components were chosen for cell clustering. Cell clusters were identified using the FindClusters function with a resolution parameter of 0.5, which employed a K-nearest neighbor (KNN) graph-based clustering approach to iteratively group cells together ^[91]^. For visualization, UMAP dimensionality reduction was performed with top 30 principal components using the RunUMAP function ^[92]^.

For cell type annotation, we detected marker genes of cell clusters by comparing each cell cluster with all other cell clusters using Wilcoxon rank sum test, with fraction of expressed cells > 0.25 in ether of the two cell populations, adjusted p-value < 0.05 and log2(fold change) > 0.25. Based on the marker genes of cell clusters, 29 major cell types were identified. Subsequently, the marker genes of cell types were detected again using the same method as described above. The complete list of marker genes for all cell types can be found in Table S3, Supporting Information.

#### 3. Projecting p-Enh-KO embryo cells onto reference map

p-Enh-KO cells were projected onto reference map by using the Seurat package. For scRNA-seq datasets from E8.5, E9.5 and E10.5 p-Enh KO embryos, gene expression normalization and variable gene identification were firstly performed using NormalizeData and FindVariableFeatures functions respectively. Next, the anchors between p-Enh-KO datasets and the reference map were identified using the FindTransferAnchors function. Finally, p-Enh-KO cells were classified and annotated using the TransferData function, as well as projected onto the reference UMAP using the MapQuery function.

#### 4. Differential expression analysis between WT and KO embryos

Comparisons between WT and p-Enh-KO were done for each cell type and at each stage. By using Wilcoxon rank sum test, differential expression genes (DEGs) were detected with fraction of expressed cells > 0.25 in ether WT or p-Enh-KO cells, adjusted p-value < 0.05 and log2(fold change) > 0.25. Enrichment analysis between DEGs and the top 100 significant marker genes of cell types were performed using Fisher’s exact test.

#### 5. Calculating module scores of a gene set at single cell levels

For a gene set of interest, the module scores in single cells were computed using the AddModuleScore function of the R package Seurat. Specially, all expressed genes were firstly divided into bins based on their averaged expression across all cells, and then control genes were randomly selected from each bin. Module scores were calculated as the difference between the average expression of the gene set of interest and the average expression of the control gene set ^[93]^.

#### 6. Analysis of anterior-posterior axis pattering and limb development

For analysis of anterior-posterior axis pattering, the cells of Forebrain/Midbrain/Hindbrain, NMP, Neural tube, Paraxial mesoderm-1, Paraxial mesoderm-2, and Somite mesoderm were selected for reclustering. For analysis of limb development, the cells of Limb mesenchyme and Mesenchyme were selected for reclustering. Consistent with the previous analysis strategy, we firstly constructed a WT reference and then projected cells of corresponding cell types from p-Enh-KO embryos onto the reference.

For RNA velocity analysis, the count matrices of pre-mature and mature mRNA abundances were firstly generated by velocyto (version 0.17.17) counting pipeline, and then used as input for scVelo (version 0.2.4) to estimate RNA velocity ^[94,95]^.

Trajectory analysis was performed using monocle3 (version 1.0.0) and pseudotime of cells was calculated based on the trajectory ^[46]^. The density plots of WT or p-Enh-KO cells along the pseudotime axis were implemented using the function kdeplot function of the python seaborn package, where a Gaussian kernel was used to produce a continuous density estimate and the density was scaled by the number of observations so as to make the total area under the density equal to 1. For posterior neural tube development and the posterior mesodermal lineages development, we selected the top 500 most variable genes, detected by using the FindVariableFeatures function of the Seurat package, as trajectory-based expressed genes.

### Gastruloids transcriptome data processing

Raw sequencing reads were firstly trimmed by Trim Galore (version 0.4.4_dev) and then aligned to mm10 reference genome by STAR (version 2.5.2b) with default parameters ^[96]^. After removal of duplicates, featureCounts (version 1.6.5) was used to count reads for genes ^[97]^. The gene annotation file (gencode.vM10.annotation.gtf) was obtained from the GENCODE project. Batch effects were corrected using the ComBat function of the sva package, where the first batch was selected as the reference batch ^[98,99]^. Differential expression analysis was performed using the R package DESeq2 (version 1.40.2) and significantly differential expression genes (DEGs) were defined as the genes with adjusted p-value less than 0.01 and fold change greater than 2 ^[100]^. GO terms enriched by DEGs was identified using the R package clusterProfiler (version 4.8.2) ^[101]^. The table containing the list of genes specifically regulated by CDX2 or p-Enh is provided in Table S5, Supporting Information.

### H3K27ac ChIP-seq data processing

Paired-end ChIP-seq reads were trimmed by TrimGalore v0.6.6 with parameters ‘-q 20’, then aligned to the mouse genome assembly (mm10) with Bowtie 2 v2.2.5 ^[102]^ and sorted with SAMtools v1.13 ^[103]^. All reads from mitochondria and chromosome Y were removed. Duplicated reads and reads with mapping quality below 30 were removed.

Peaks were called by MACS2 v2.2.7.1 ^[84]^ with parameters ‘--extsize 200 --nomodel -- nolambda --keep-dup all -p 0.01 -g mm’. Differential peak analysis was performed with the DESeq2 method in DiffBind 3.8.4 ^[104]^. Significant differential peaks were defined with FDR <0.05 and log2(FoldChange)>=1. Bigwig files were normalized in RPKM under 50bp bin and generated with deepTools v3.5.1 bamCoverage ^[105]^. Annotations of peaks were performed with R package ChIPSeeker v1.5.1 ^[106]^. Table containing detailed information about differential H3K27ac ChIP-seq peaks is provided in Table S6, Supporting Information.

### TFs ChIP-seq data processing

TFs ChIP-seq includes SMAD2/3/4, CDX2 from different public datasets with paired-end or single-end sequencing. Paired-end and single-end reads were trimmed by TrimGalore v0.6.6 with parameters’-q 20’ relatively, then aligned to the mouse genome assembly (mm10) with Bowtie 2 v2.2.5 ^[102]^ and sorted with SAMtools v1.13 ^[103]^. All reads from mitochondria and chromosome Y were removed. Duplicated reads and reads with mapping quality below 30 were removed.

Peaks were called by MACS2 v2.2.7.1^[84]^ with parameters ‘--extsize 200 --nomodel --keep-dup all -p 1e-5 -f BAM -g mm’. Bigwig files were normalized in RPKM under 50bp/10bp bin and generated with deepTools v3.5.1 ^[105]^ bamCoverage.

### Signaling pathway enrichment analysis

GEO-seq data for E7.5 mouse embryo were obtained from the NCBI GEO under accession numbers GSE120963 ^[9]^. Processing of GEO-seq data samples started with raw sequencing reads. Quality control metrics such as sequence quality scores, GC content, and adapter content were assessed using FastQC (v0.11.9). Low-quality reads and adapters were trimmed or removed. Then, the high-quality reads were aligned to a reference genome (mm10) with Hisat2 (v2.2.1) ^[107]^. Gene expression quantification was performed using featureCounts (v1.5.3) ^[97]^. The count matrix generated representing the number of reads mapped to each gene in each sample. The count matrix was normalized to account for differences in sequencing depth and gene length as Transcripts Per Kilobase of exon model per Million mapped reads (TPM). To evaluate the enrichment of genes response to TGF-β, AUCell (v1.20.2) was used to calculate the gene-set activity score (gene set from GESA: MM9770) ^[108]^. The score was calculated by the top 10% genes of all genes in expression matrix.

## Supporting Information

Figure S1-8; Table S1-6

## Acknowledgments

We are grateful to Prof. Chi-Chung Hui for his constructive suggestions. We thank the core facilities at Guangzhou National Laboratory and Center for Excellence in Molecular Cell Science, CAS for their excellent support, especially in animal care and imaging support.

## Author Contributions

Conceptualization: X.Y, N.J., Y.C.

Investigation: Y.C., F.T., X.Y., L.Z., Q.F., J.L., P.S., M.W., Y.Q., R.S., Y.F., H.J.X., R.W.

Visualization: Y.C., X.Y., F.T., Q.F., R.S., Y.F.

Funding acquisition: X.Y., N.J.

Supervision: X.Y., N.J., J.L., Z.S., C.L.

Writing – original draft: Y.C., X.Y., F.T., Q.F., N.J.

Writing – review & editing: X.Y., N.J., Y.C., F.T., Q.F., C.L.

## Data and materials availability

All raw and processed sequence data reported in this paper have been deposited in the Genome Sequence Archive ^[109]^ in National Genomics Data Center ^[110]^, China National Center for Bioinformation / Beijing Institute of Genomics, Chinese Academy of Sciences (GSA: CRA014616) that are publicly accessible at https://ngdc.cncb.ac.cn/gsa.

