## Supplementary Figures for "Gastrula-premarked posterior enhancer primes posterior tissue development through cross-talk with TGF-β signaling pathway"

### Figure S1. Screening and identification of posterior development related distal regulatory elements

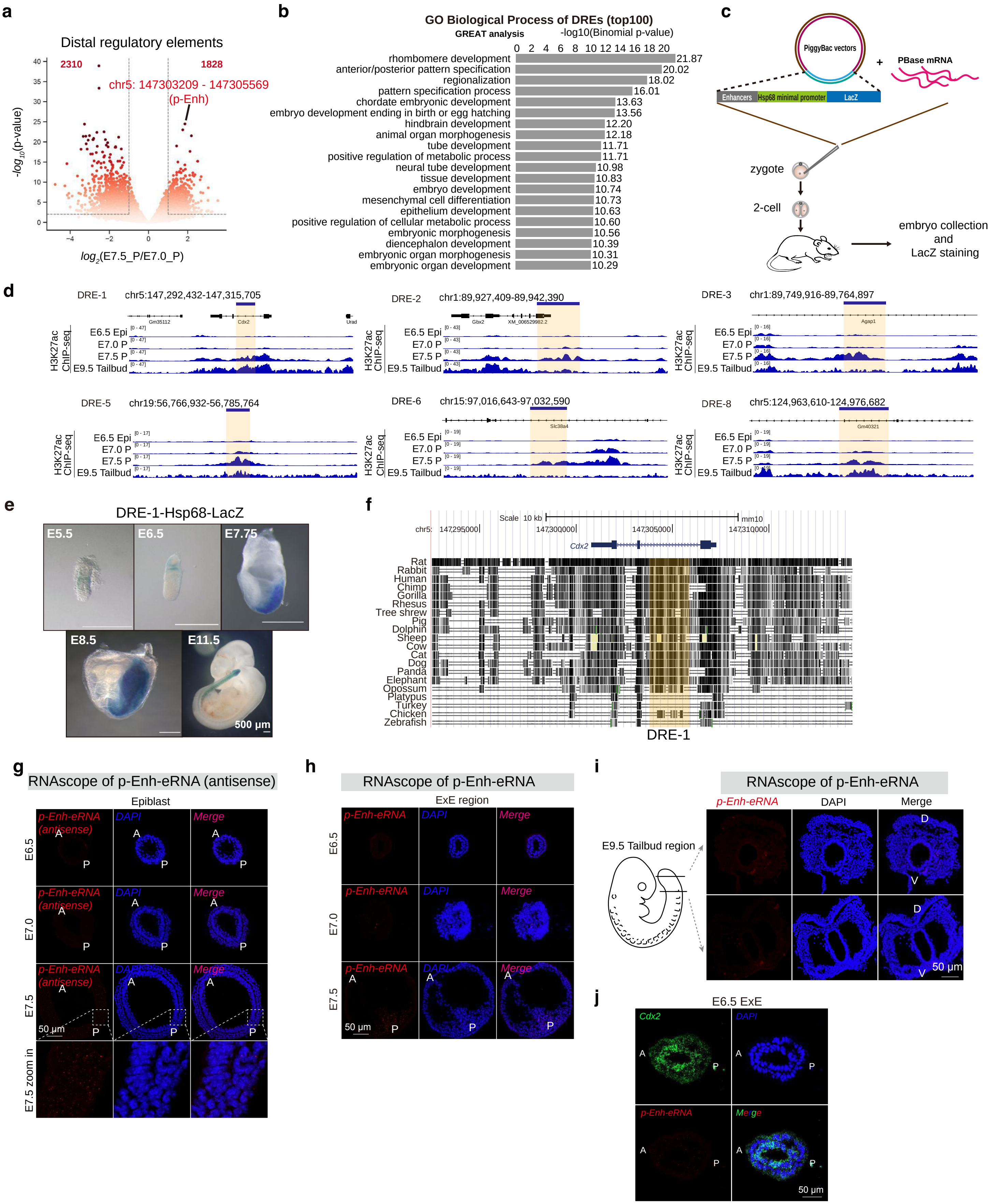

**Figure S1. Screening and identification of posterior development related distal regulatory elements.**

**(a)** Volcano plot illustrating the posterior-specific distal regulatory elements with significant changes in activity.

**(b)** GO biological process enrichment analysis (GREAT) of top-100 DREs. Detailed information regarding the top-100 DREs is provided in Table S1, Supporting Information.

**(c)** The diagram illustrating the experiment details about the transgenic embryo screening system for analyzing the activity of DREs during embryonic development.

**(d)** IGV snapshots of H3K27ac ChIP-seq signals of indicating DREs (correspond to Figure 1d).

**(e)** DRE-1-Hsp68-LacZ transgenic embryos illustrating robust activity of p-Enh throughout various developmental stages. Scale bar: 500 μm.

**(f)** UCSC browser snapshot showing high conservation of DRE-1 across mammalian species.

**(g)** RNAscope results of antisense p-Enh-eRNA in epiblast regions from E6.5 to E7.5. A: anterior region, P: posterior region. Scale bar: 50 μm.

**(h)** RNAscope results of p-Enh-eRNA in ExE regions from E6.5 to E7.5. A: anterior region, P: posterior region. Scale bar: 50 μm.

**(i)** RNAscope targeting p-Enh-eRNA in E9.5 tailbud regions. D: dorsal region, V: ventral region. Scale bar: 50 μm.

**(j)** Co-staining of *Cdx2* mRNA and p-Enh-eRNA using RNAscope in the ExE regions in E6.5 mouse embryos. Scale bar: 50 μm.

### Figure S2. p-Enh-KO mouse embryos exhibit embryonic lethality

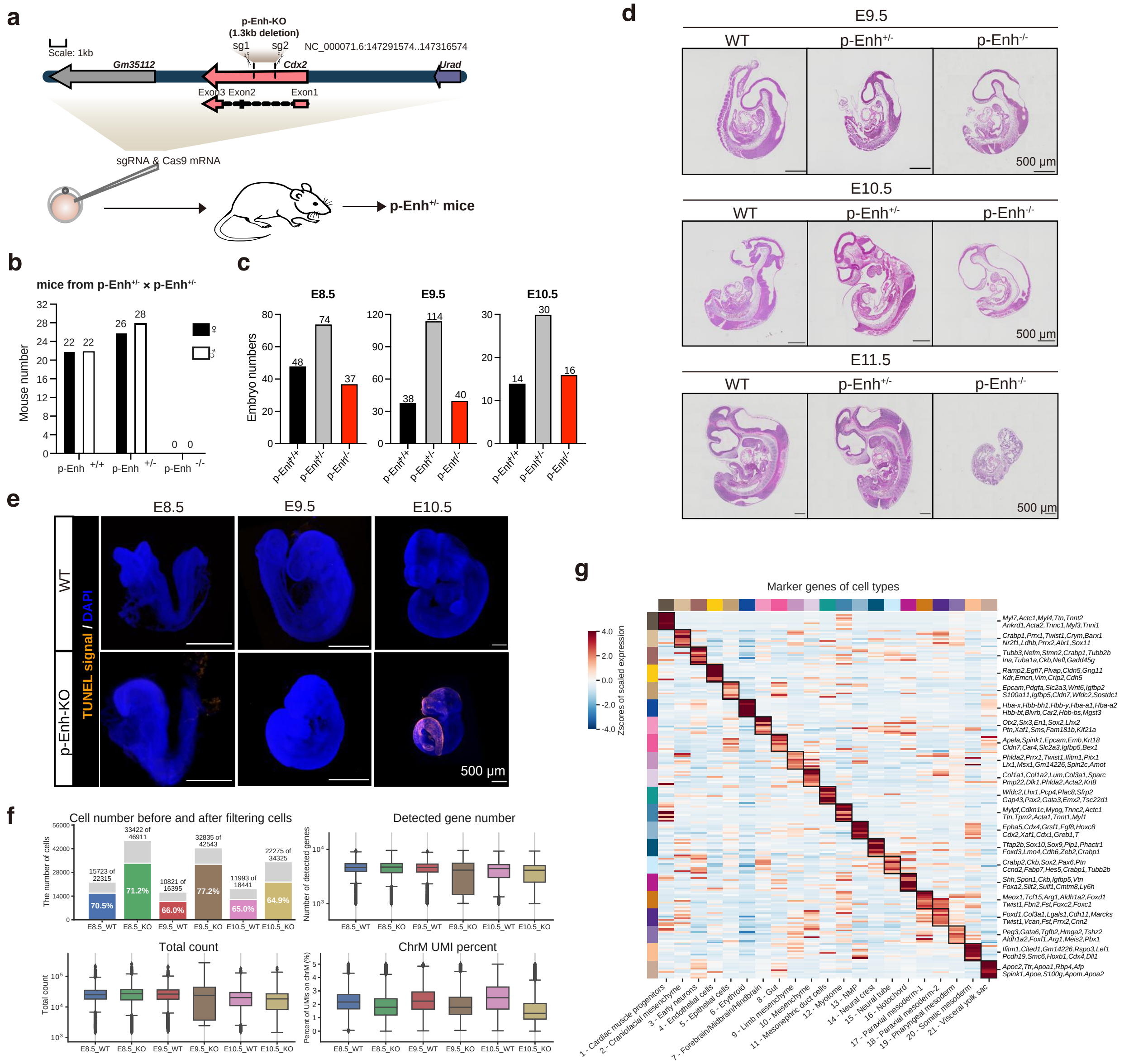

**Figure S2. p-Enh-KO mouse embryos exhibit embryonic lethality.**

**(a)** The diagram illustrating the strategy for generating p-Enh-KO embryos. Two pairs of sgRNA targeting boundaries of p-Enh locus were shown.

**(b)** Genotypes of offspring from self-crossing of p-Enh$_{+/-}$ mice.

**(c)** Genotypes of embryos at different developmental stages from self-crossing of p-Enh$_{+/-}$ mice.

**(d)** HE staining results of embryos with different genotypes at different developmental stages. Scale bar: 500 μm.

**(e)** Lightsheet imaging of whole-mount TUNEL results of WT and p-Enh-KO embryos at different development stages. Scale bar: 500 μm.

**(f)** Quality control of single-cell RNA-seq data.

**(g)** Heatmap showing top-10 marker genes of each cluster. The complete list of marker genes for all clusters can be found in Table S3, Supporting Information.

### Figure S3. Developmental deficiency in p-Enh-KO embryos

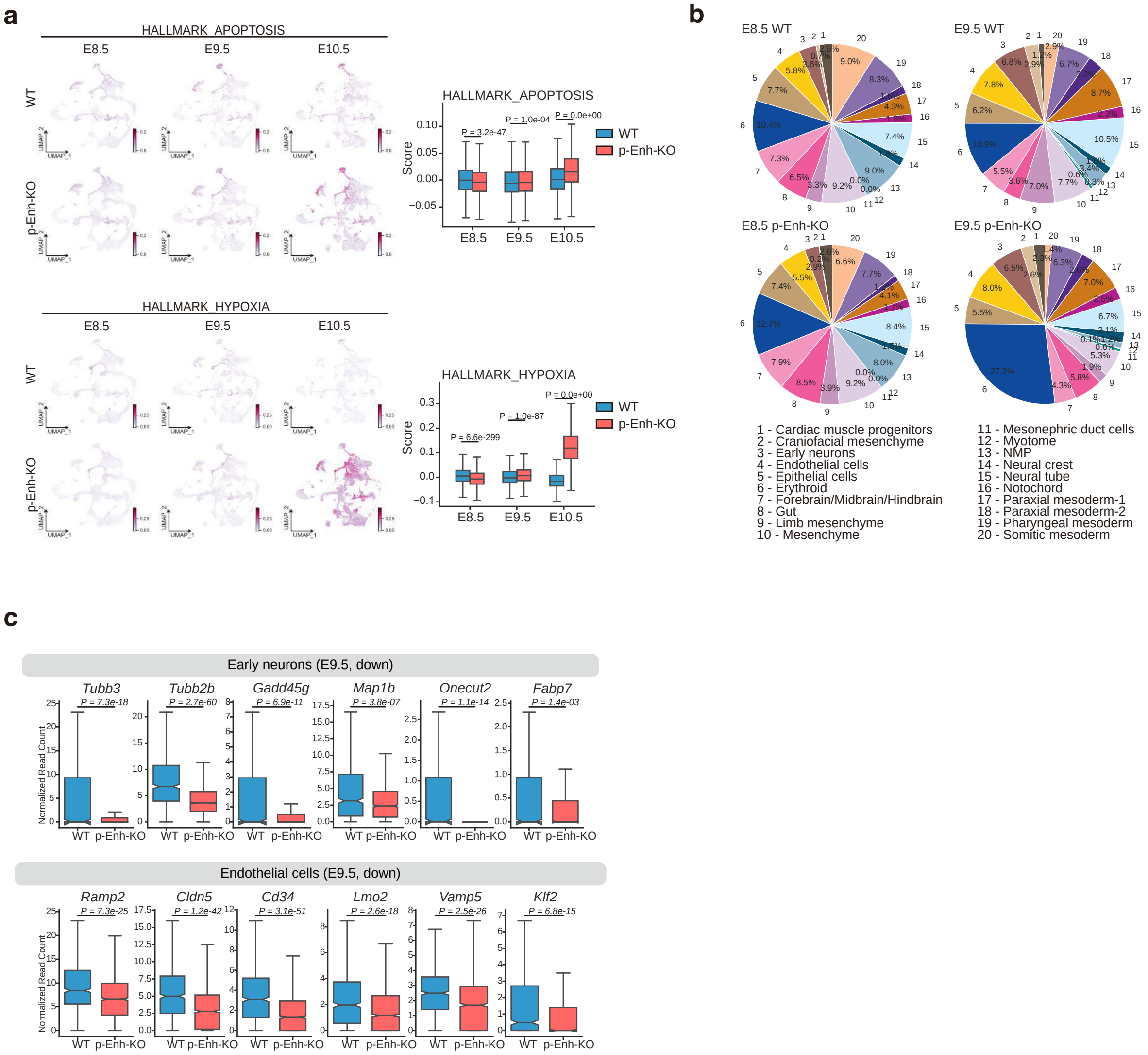

**Figure S3. Developmental deficiency in p-Enh-KO embryos.**

**(a)** Activities of apoptosis and hypoxia signaling were assigned based on hallmark gene list from GSEA dataset. Boxplot illustrating the corresponding activity score in each sample. p-values were labeled on the boxplot.

**(b)** Percentage of cell abundance of different clusters in each sample.

**(c)** Boxplot illustrating normalized read count of marker genes with significant expression changes between WT and p-Enh-KO embryos in indicated clusters. p-value was indicated in each gene.

### Figure S4. Reconstruction of digital A-P axis

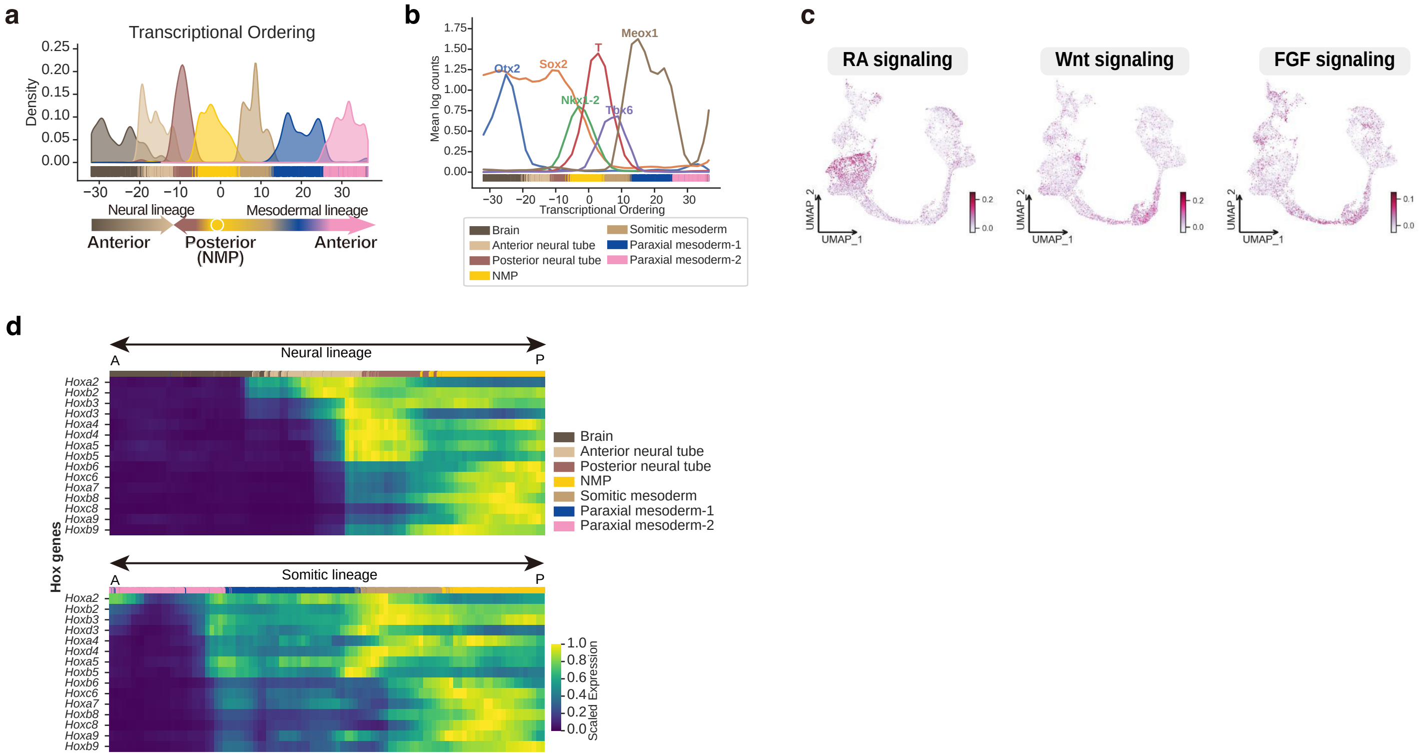

**Figure S4. Reconstruction of digital A-P axis.**

**(a)** Cell density distribution along the transcriptional ordering. Two different origins of neural tube cells were indicated based on RNA velocity result in Figure 3a.

**(b)** The expression patterns of selected transcription factors along the one-dimensional A-P axis.

**(c)** RA, Wnt, and FGF signaling activities projected along the reconstructed A-P axis UMAP. Signaling activities were measured through gene sets (MM15186, MM3864, MM5178) from GSEA datasets [78].

**(d)** Heatmap showing the expression patterns of *Hox* gene family along the one-dimensional transcriptional ordered A-P axis.

### Figure S5. Generation of p-Enh-KO and Cdx2-KO mESC and their differentiation deficiency

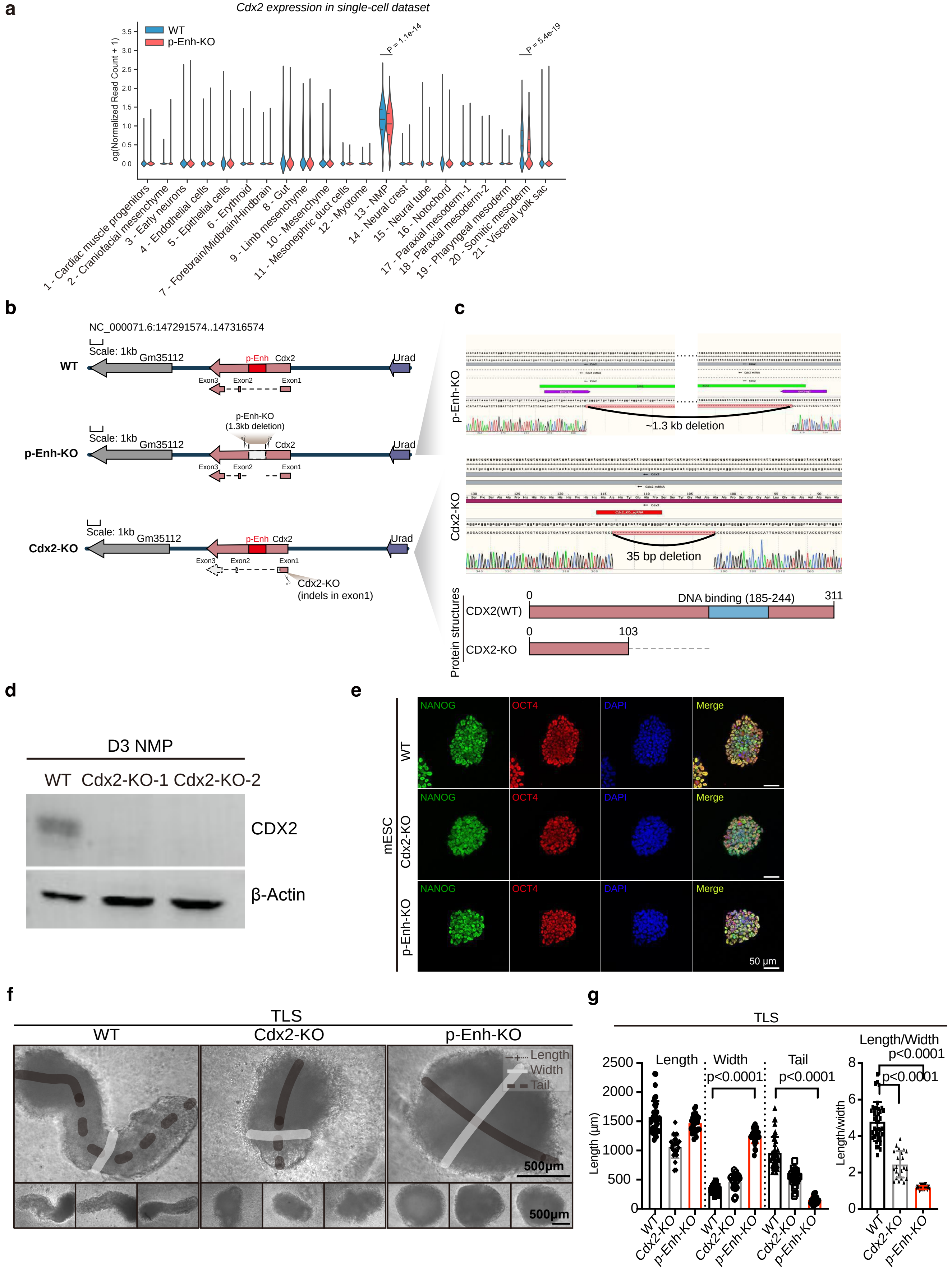

**Figure S5. Generation of p-Enh-KO and Cdx2-KO mESC and their differentiation deficiency.**

**(a)** The expression pattern of *Cdx2* in different cell clusters in single-cell dataset of WT and p-Enh-KO embryos.

**(b)** Diagram illustrating the strategy for generating p-Enh-KO and Cdx2-KO mESC.

**(c)** Sanger sequencing results showing successfully knock out of p-Enh-KO and induced indel in Cdx2-KO cell lines. The protein structure diagram showing the absence of CDX2 DNA binding domain in Cdx2-KO cellline.

**(d)** Western blotting result for CDX2 protein in NMP cells.

**(e)** Immunofluorescence results of store cultured mESC of WT, Cdx2-KO and p-Enh-KO cell lines. Scale bar: 50 μm.

**(f)** Bright field images of TLS with distinct genotypes.

**(g)** Statistical analysis of morphology parameters of TLS in each group.

**Figure S6. Transcriptome analysis revealed distinct functions between p-Enh and *Cdx2***

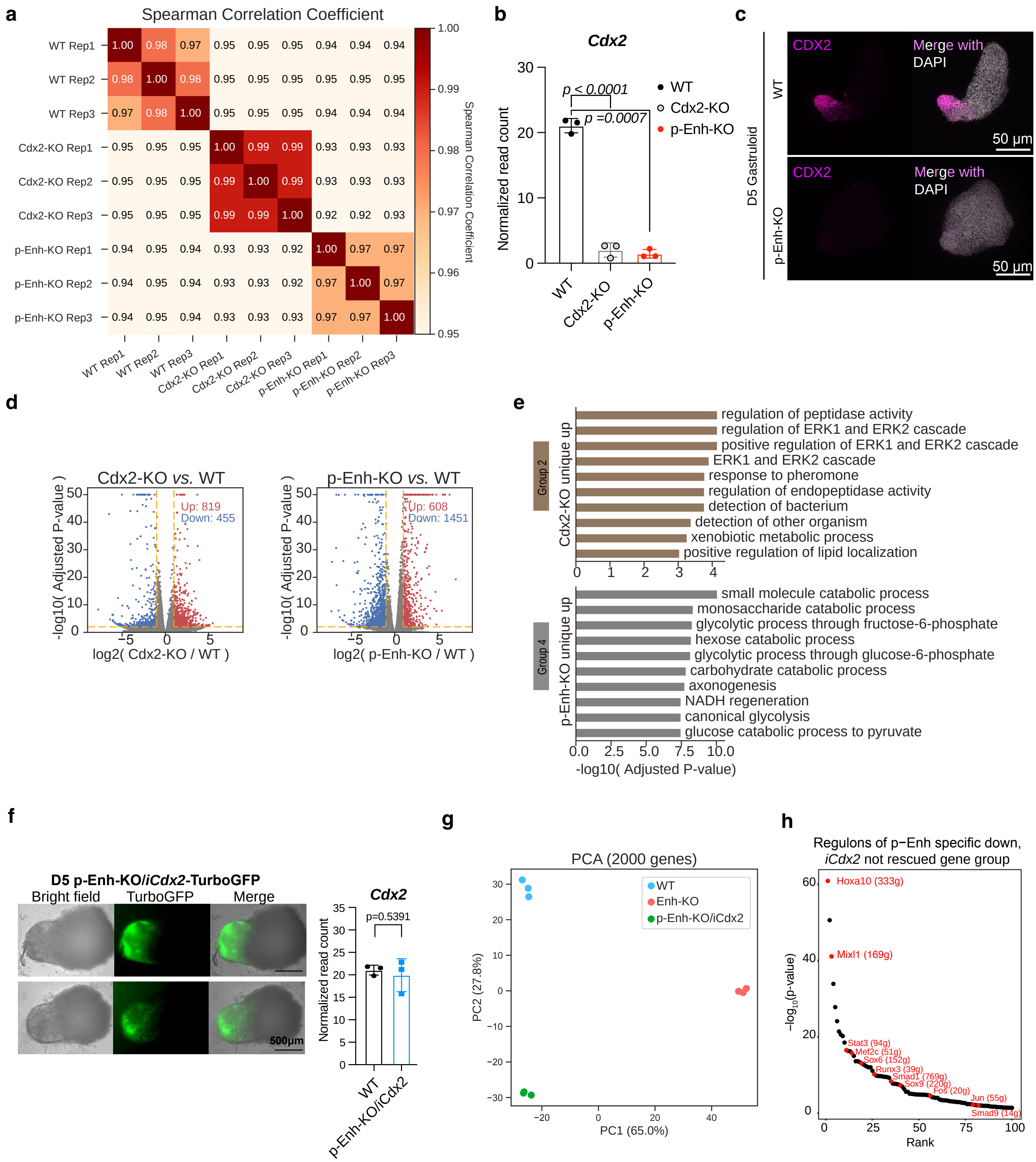

**Figure S6. Transcriptome analysis revealed distinct functions between p-Enh and *Cdx2*.**

**(a)** Heatmap showing spearman correlation coefficient among different transcriptome data from WT, Cdx2-KO and p-Enh-KO groups.

**(b)** Normalized read count of *Cdx2* in different gastruloid samples.

**(c)** iDISCO immunofluorescence results targeting CDX2 protein in D5 gastruloids from WT and p-Enh-KO groups. Scale bar: 50 μm.

**(d)** Differential expressed genes (DEGs) (p-value < 0.01, fold change >2) of Cdx2-KO vs. WT, and p-Enh-KO vs. WT. The up-regulated genes were labeled in red and the down-regulated labeled in blue.

**(e)** Gene Ontology (BP) results of Group 2 and Group 4 genes defined from Figure 4h.

**(f)** Bright field image and TurboGFP expression in D5 p-Enh-KO/iCdx2 gastruloids. Normalized read count of *Cdx2* transcripts confirmed successful overexpression.

**(g)** Principal component analysis result of gastruloid transcriptomes from WT, p-Enh-KO, and p-Enh-KO/iCdx2 groups.

**(h)** Regulons of p-Enh-KO specific down, *iCdx2* could not rescued gene groups (Group 5-B). Regulons overlapped with TGF-β signaling pathway were highlighted in red.

### Figure S7. Loss of p-Enh leads to genome-wide epigenomic remodeling

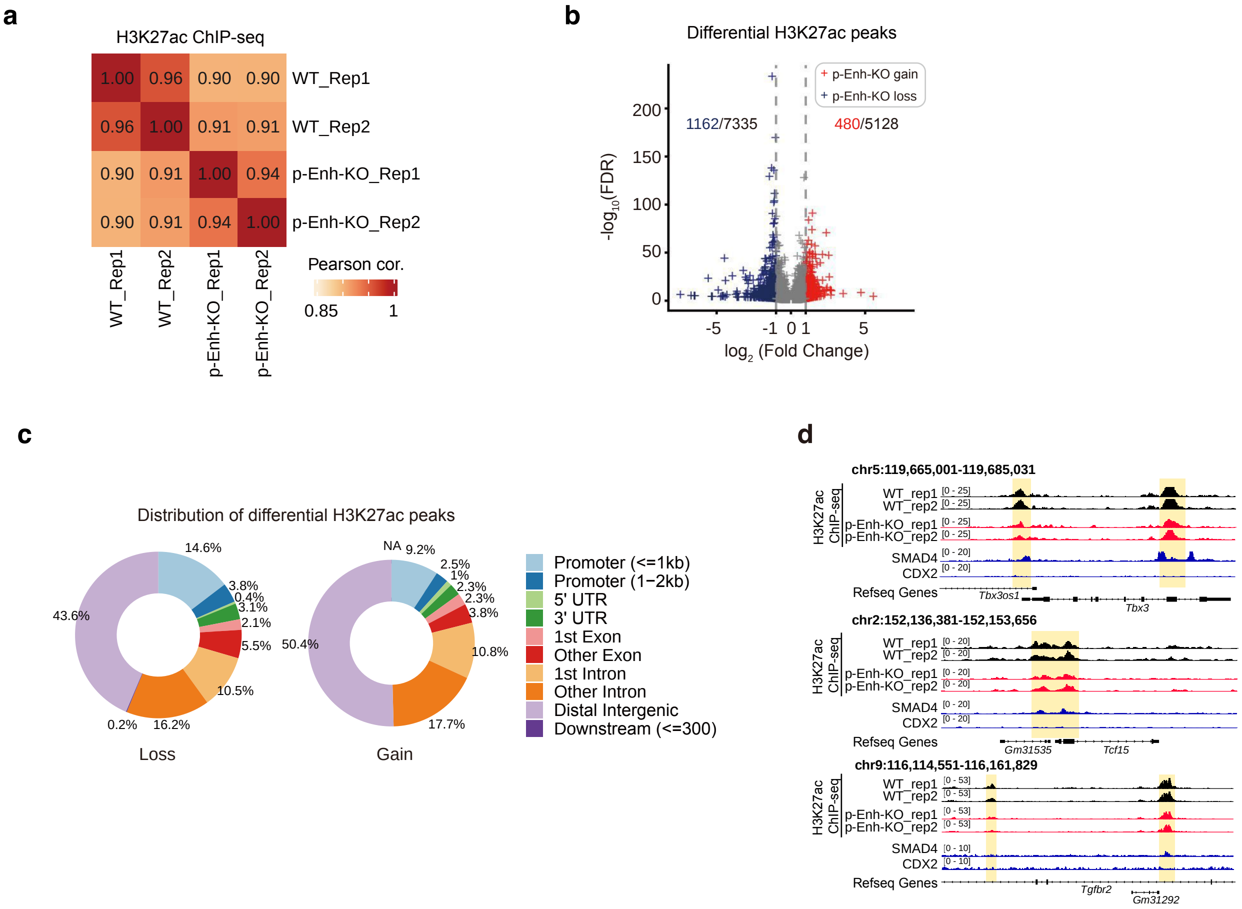

**Figure S7. Loss of p-Enh leads to genome-wide epigenomic remodeling.**

**(a)** Heatmap showing Pearson Correlation Coefficient among H3K27ac ChIP-seq data from WT and p-Enh-KO NMPs.

**(b)** Volcano plot illustrating differential H3K27ac peaks. Significant differential peaks were defined with FDR <0.05 and log2(Fold Change)>=1.

**(c)** Distribution of p-Enh-KO lost and gained peaks.

**(d)** IGV snapshots of representative loci with obvious alterations in chromatin activities, which exhibit overlap with the binding peaks for SMAD4, rather than CDX2.

### Figure S8. Transcriptome analysis of the effect of Activin A in gastruloid system

**a**

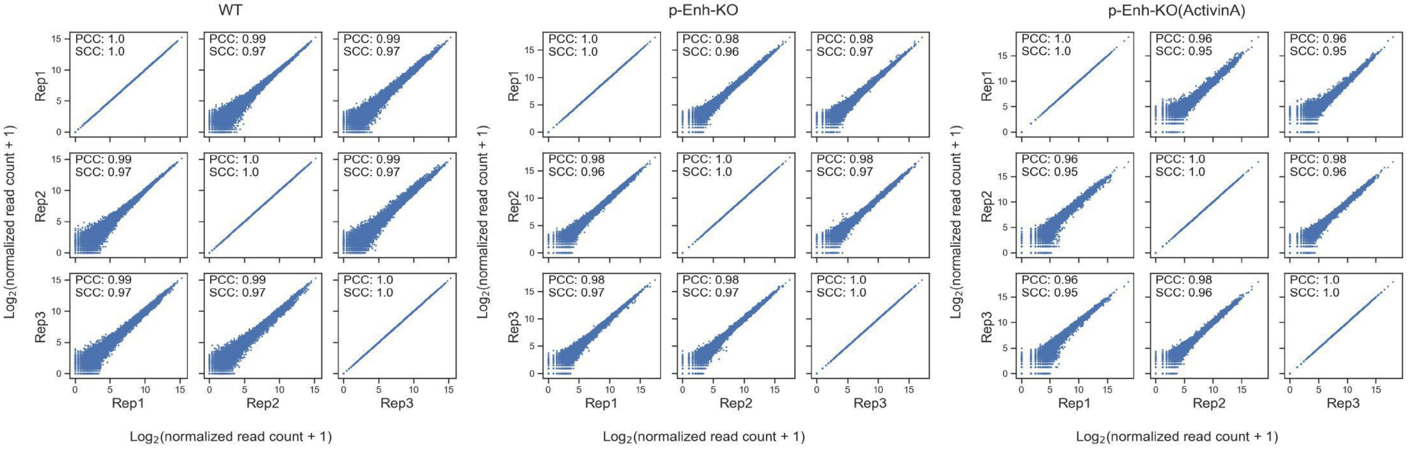


**b**

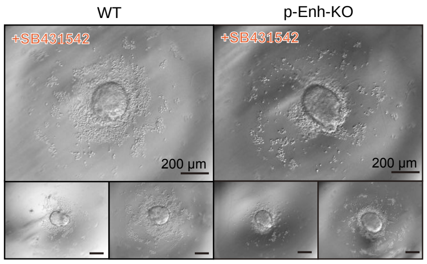

**Figure S8. Transcriptome analysis of the effect of Activin A in gastruloid system.**

**(a)** Scatter plots showing correlation among transcriptome data from WT, p-Enh-KO, p-Enh-eRNA-KD, and p-Enh-KO (ActivinA) groups.

**(b)** Bright field images of D5 gastruloids from WT and p-Enh-KO groups in SB431542 treatment condition.
